# Cross-platform digital PCR evaluation of bovine papilloma virus quantification: introducing PCR-ValiPal for Standardized guided assay validation

**DOI:** 10.1101/2025.09.04.674204

**Authors:** David Gleerup, Matthijs Vynck, Lien Gysens, Cindy De Baere, Jo Vandesompele, Olivier Thas, Ann Martens, Maarten Haspeslagh, Ward De Spiegelaere

## Abstract

Digital Polymerase chain reaction (dPCR) enables precise and absolute quantification of nucleic acids by partitioning samples into thousands of individual PCR micro-reactions. While it minimizes the need for standard curves and enhances reproducibility compared to qPCR, thorough assay validation remains crucial. We introduce PCR-ValiPal, a user-friendly web application that standardizes dPCR assay validation steps and streamlines calculations of limit of blank (LOB), limit of detection (LOD), limit of quantification (LOQ), precision, trueness, and linearity in accordance with International Organization for Standardization (ISO) 20395:2019. To demonstrate PCR-ValiPal’s capabilities and the value of method-specific optimization, we use it to validate a novel three-color PCR assay for Bovine Papillomavirus (BPV) types 1 and 2, comparing four platforms: Naica (droplet dPCR), QIAcuity (microwell dPCR), LOAA (real-time dPCR), and CFX96 (qPCR). Using synthetic standards, we assess the platforms’ performance under identical assay conditions. Naica and QIAcuity showed lower LOB and LOQ values, along with minimal bias for BPV-1, while LOAA demonstrated stable but negative bias. Although qPCR exhibited the highest sensitivity for BPV-2, it was less sensitive at low concentrations for BPV-1. These results underscore the value of method-specific optimization and the usefulness of PCR-ValiPal.

## Introduction

Digital PCR (dPCR) offers highly accurate and absolute quantification of nucleic acids, enabling a broad range of applications in areas such as microbiology (1, 2), oncology (3, 4), copy number variation (5, 6), and environmental DNA surveillance (7, 8). By partitioning each nucleic acid sample together with PCR reagents into microwells or droplets, dPCR creates thousands of parallelized PCR reaction containers, called partitions, and allows for the direct counting of the presence or absence of target molecules. This absolute quantification improves precision, reduces reliance on certified reference materials, and enhances intermediate precision (9–11). Despite these strengths, rigorous assay validation remains crucial for ensuring reliable performance, including the determination of limit of blank (LOB), limit of detection (LOD), limit of quantification (LOQ), repeatability, and intermediate precision (Table 1).

**Table 1.**
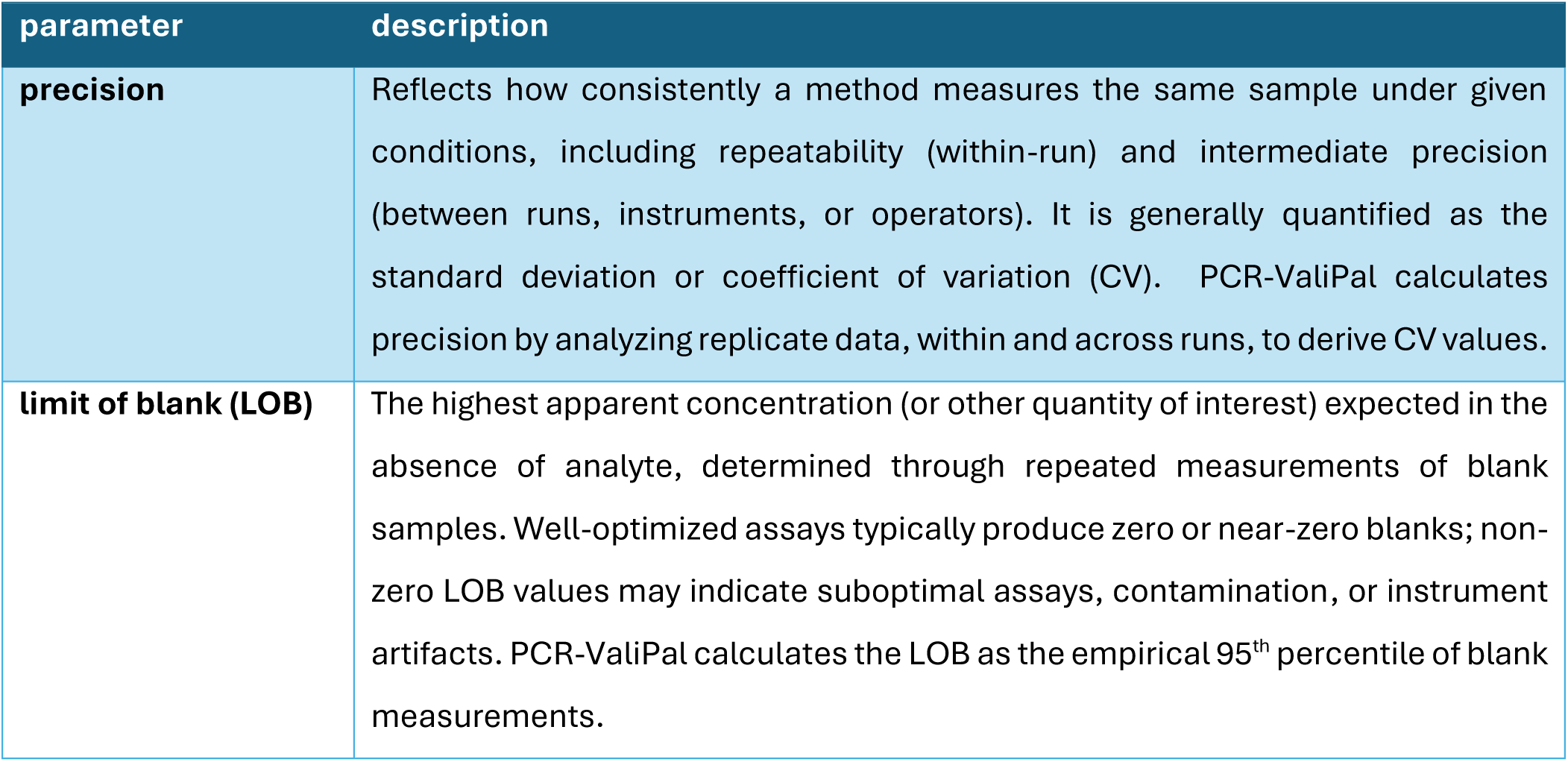
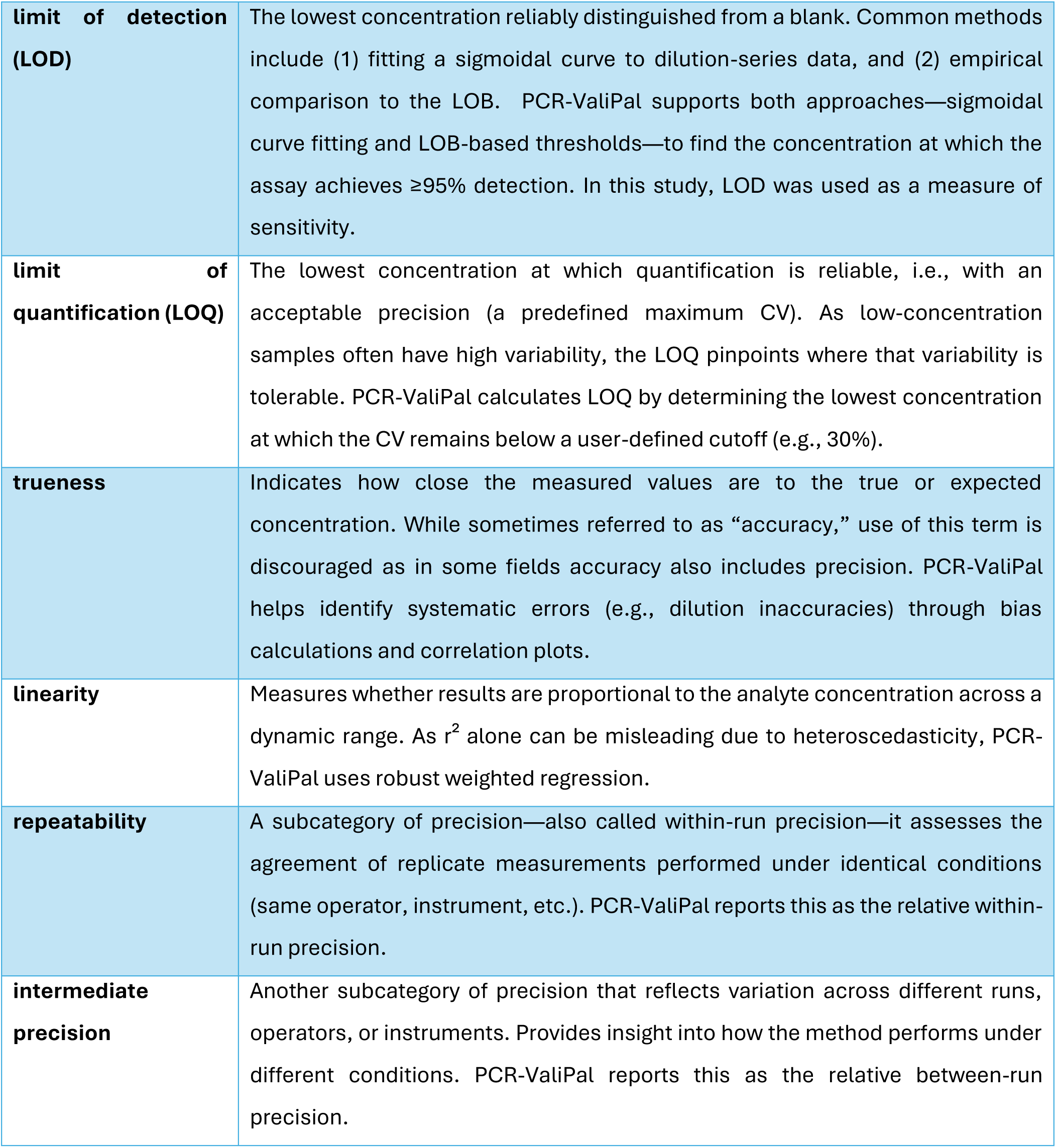
Expanded analytical parameters and definitions (based on ISO 20395:2019 and further guidance from Vynck et al., 2017 and Deprez et al., 2016 (14–16)).

Although several guidelines, e.g., MIQE for qPCR (12), dMIQE for dPCR (13) and ISO20395:2019 for both qPCR and dPCR (14), emphasize detailed reporting and validation, many of these steps are laborious, both in the wet lab and during data analysis. Yet, such reporting and validation are essential for ensuring reliability. To facilitate and standardize this process – also for the non-expert - we developed PCR-ValiPal. This web tool is a rigorous update of the previously published dPCalibRate and calculates the comprehensive set of analytical parameters required for assay validation and reporting under ISO20395:2019 requirements (Table 1) (15).

To showcase an ISO-guided assay validation report using PCR-ValiPal, a technical cross-platform dPCR validation study was performed using a three-color dPCR assay for the detection of several bovine papilloma virus (BPV) strains using synthetic templates.

Bovine papillomavirus type 1 (BPV-1) is a double-stranded DNA virus from the *papillomaviridae* family that primarily infects cattle, leading to papillomas in epithelial and fibroblastic tissues. Notably, BPV-1 can induce tumorigenesis in its natural bovine hosts and, experimentally, in other species such as horses, where it causes equine sarcoids (17). As such, BPV-1 serves as a valuable model for studying papillomavirus biology and cancer development.

While BPV-1 is the predominant type detected in equine sarcoids in Europe (18), BPV-2 has also been identified in European equids, albeit less frequently (19). Additionally, co-infections with BPV-1 and BPV-2 have been documented. A study analyzing 104 equine sarcoid samples from New Zealand reported that approximately 10% of the lesions contained both BPV-1 and BPV-2 DNA (20).

Therefore, the BPV assay was designed to detect both BPV-1 and BPV-2 strains, as well as the INF reference gene target for normalization purposes. This comprehensive detection is valuable for both epidemiological surveillance and clinical outcome studies, as understanding the distribution and co-infection dynamics of BPV strains can inform disease management and treatment strategies.

## Materials and methods

### Data analysis using PCR-ValiPal

#### Limit of blank

The limit of blank (LOB) is often defined as

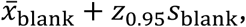

with *x̅*_blank_ the mean concentration observed in the blank samples, *s*_blank_ the standard deviation of the blank samples, and *z*_0.95_ the 95% quantile of the standard normal distribution.

This definition assumes a normal distribution of concentrations. This assumption is often violated: for a well-optimized assay, the majority of the negative control reactions are observed with a concentration of zero. PCR-ValiPal implements an alternative that estimates the empirical 95% quantile of all observed concentrations in the blank samples. This avoids the assumption of a normal distribution of concentrations and is more appropriate for PCR experiments. A drawback of this approach is that it requires many blank samples to be analyzed to obtain a reliable estimate of the 95% quantile.

In this context, LOB samples refer to negative controls — samples containing the sample matrix, in this case background DNA, but not the target. These should not be confused with non-template controls that contains water instead of template. In general 60 LOB control replicates are recommended for validating a novel assay, but 20 LOB replicates are deemed sufficient for verifying the LOB of an established assay (21). Note also that that these should be assessed using the complete assay to account for any potential probe or primer interactions. Furthermore, LOB replicates should not be tested sequentially, but should be interspersed between the samples (14).

#### Limit of detection

The limit of detection (LOD) is the lowest concentration reliably distinguished from the LOB. PCR-ValiPal implements two distinct approaches for estimating the LOD:

Type 1 LOD: positive sample approach

∘ The LOD is the highest dilution level (lowest concentration) at which a user defined percentage (by default 95%) of replicate measurements exceed the LOB (21, 22).
∘ This method may be more variable with a low number of technical replicates but takes the LOB into account and may return more appropriate values when the LOB is not zero.

Type 2 LOD: Sigmoidal curve approach

∘ A logistic curve is fit to the fraction of positive reactions at each dilution level. The LOD is the concentration observed where the curve intersects the chosen detection probability *β* (e.g., 95%) (23).
∘ Optionally, a bootstrap method generates a confidence interval by resampling the observed fraction of positives at each of the dilution levels and re-fitting the logistic curve. For each of 1000 bootstrap resamples, an LOD can be calculated as before. A 1 − *α*% confidence interval is then obtained by calculating the *α*/2 and 1 − *α*/2 quantiles of the LODs.

Note, in this context, LOD determination requires replicate measurements close to the expected detection limit. In line with CLSI EP17-A recommendations, at least 20 replicate measurements should be obtained to ensure reliable estimation(22).

#### Limit of quantification

The LOQ identifies the lowest concentration at which the assay maintains an acceptable level of precision. PCR-ValiPal determines two types of LOQ. The first is defined by comparing the coefficient of variation (CV) to a user-defined cutoff (default: 30%) and identifying the lowest concentration where the CV is below the cutoff, provided that all higher concentrations also remain below the cutoff. The second is obtained similarly but uses the CV’s upper 95% confidence instead of the CV itself when compared against that user-defined cutoff and is a more conservative approach.

#### Trueness

Trueness describes how close measured concentrations are to the true (expected) value, for a given dilution level. In PCR-ValiPal, bias is estimated as

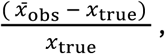

with *x̅*_obs_ the mean observed concentration, and *x*_true_ the expected concentration. A positive bias indicates overestimation, whereas a negative bias indicates underestimation of the expected concentration.

#### Linearity

Linearity assesses whether the measured concentration scales proportionally over a range of analyte levels. PCR-ValiPal fits a robust weighted least squares (WLS) model and then tests whether the quadratic term is significantly different from zero, indicating non-linearity. While often used to claim linearity, a high r² in isolation can be misleading, in particular when the variance changes with the expected concentration (heteroscedasticity). Therefore, a statistical verification of non-linearity using said quadratic regression is recommended.

#### Repeatability and Intermediate Precision

To evaluate the within-run repeatability and between-run intermediate precision according to ISO 20395:2019 (14), PCR-ValiPal employs an ANOVA-based variance component analysis. Briefly, each dilution level (or sample) is measured in multiple replicates per run, with the total number of runs spanning multiple days and/or operators. PCR-ValiPal then partitions the overall variance into the within-run variance (repeatability) and the between-run variance (intermediate precision; Deprez et al. 2016 (16)).

All calculations in PCR-ValiPal align with the ISO 20395:2019 guidelines and standard practices (15, 16). By integrating these statistical analyses in a single tool, users can easily estimate the precision, LOB, LOD, LOQ, trueness, and linearity—facilitating transparent and reproducible assay validation.

### Experimental methods and design

#### Synthetic template

As positive control material for the primer/probe assays, 126 bp gBlock Gene Fragments (Integrated DNA Technologies, Coralville, IA, USA) were synthesized based on the amplicons generated by the BPV-1, BPV-2 and IFN assays (see section S.1 for gBlock sequences). Each gBlock was resuspended in TE buffer (10 mM Tris, 1 mM EDTA) to 10 ng/µL. They were initially diluted in 1:100 steps, then mixed to create triple positive sample material, resulting in a further 1:3 dilution and a stock concentration of 1.2 million DNA copies per 1.5 µL. This stock was aliquoted for use on each platform and stored at −20 °C, then measured in duplicate on the Naica system using the protocol described below. Differences in starting concentration were observed between aliquots; however, the previously determined dilution steps were followed for each system, albeit resulting in slightly different expected values. Each aliquot was used to prepare a serial dilution on the day of measurement, with further aliquots made to maintain the same freeze/thaw cycle. All dilutions were performed in TE + background DNA (bovine genomic DNA, muscle tissue origin, 20 ng/µL), serving both as carrier nucleic acid and to approximate the background matrix. The triple positive stock yielded a 17-point dilution series (expected values in Supplementary Table S.2; corresponding dilution points in Supplementary Table S.3). For dPCR systems, only 13 points were used, due to dynamic range considerations and the limited number of wells while maintaining the minimum number of replicates per day. The concentration range spanned from 1 × 10⁶ to 0.625 DNA copies per 1.5 µL, the sample input volume for each platform. A constant input volume of 1.5 µL was maintained across platforms to enable direct comparison of assay performance. This choice reflects the volume used in qPCR for this assay in clinical practice and was therefore considered representative of the intended application rather than of each platform’s theoretical maximum sensitivity. Blank samples consisted of 1.5 µL of TE with background DNA but without synthetic oligos.

#### Experimental design

To assess the assay metrics across the different platforms at a level close to that recommended in ISO 20395:2019, minor deviations were made due to the maximum number of wells available per run on the dPCR systems. Specifically, most measurements were performed in nine replicates (with ISO recommending ten), while an additional replicate was included for concentrations around the expected LOD, resulting in ten replicates at those levels. The gBlock dilution series was analysed in triplicates, each day, on 3 different days, with an extra replicate around the expected LOD for a total of 9-10 replicates per concentration level. To include inter-operator variability in the downstream calculations, each dilution series was made by a different person. To measure the LOB on each system, 20 NTCs were analysed on each dPCR system (Naica, QIAcuity and LOAA, see further for details), while 60 were analysed on the qPCR instrument (Bio-Rad CFX96, see further for details). Of note, the qPCR instrument is located in a different laboratory than the dPCR platforms, as such, there are some differences in the reagents used in the protocols (e.g., distilled water for qPCR versus HPLC water for dPCR). In compliance with dMIQE guidelines 1-color plots from all platforms have been included in supplementary figures S.1 – S.8. LOAA BPV-1 and BPV-2 was included as a 2-color plot as it was the only option. Additionally, average lambda values with standard deviation and average partition count with standard deviation is reported in tables S.4 and S.5 respectively. Finally, all PCR-ValiPal input data is included in supplementary tables S.6 – S.17 found in the supplementary data file.

#### Bio-Rad quantitative PCR

The qPCR runs were performed in a Bio-Rad CFX96 Real-Time System (Bio-Rad, Hercules, CA, USA) in a final volume of 15 μL reaction mixture containing 0.33 µM of all primers (BPV-1, BPV-2 and IFN), 1 µM of each probe, 1X iQ Supermix (Bio-Rad), 1.5 μL of DNA and final volume made up with distilled water. Cycling conditions were 95 °C for 3 min, then 40 cycles of 95 °C for 15 s and 57 °C for 30 s. A calibration curve was made using the approach described in the synthetic template material section and served as the basis for quantifying the other dilution series.

#### Stilla Naica Droplet dPCR

The dPCR assay was carried out on the Naica dPCR system (Stilla Technologies, Villejuif, France) in a final volume of 8 µL reaction using Opal chips. The reaction consisted of 1X PerFecTa Multiplex qPCR ToughMix, 1 µM primers, 0.25 µM probes and 1.5 µL of DNA and 1 µM fluorescein, the final volume made up with HPLC grade water. Cycling conditions were 95 °C for 3 min, then 40 cycles of 95 °C for 15 s and 57 °C for 30 s, afterward an additional 5 cycles of 95 °C for 15 s and 55 °C for 30 s were used to boost fluorescence intensity as described (24). Thresholding was done manually, and a unique threshold was applied to each well due to observed baseline shifts between wells. Analysis was done using Crystal Miner software version 4.

#### Qiagen Qiacuity microwell dPCR

The microwell dPCR was done using the Qiacuity dPCR system (Qiagen, Hilden, Germany) using the 26k 24-well nanoplates with a final reaction volume of 40 µL, the final mix consisted of all primers at 0.8 µM and all probes at 0.4 µM, QIAcuity probe PCR kit at 1X and 1.5 µL of DNA. The remaining volume was HPLC grade water. The cycling conditions were as follows: 2 min at 95 °C followed by 40 cycles of 95 °C for 15 s and 57 °C for 30 s. Thresholding on the QIAcuity mainly used the built-in algorithm of QIAcuity software suite 1.2, this was done because there was a good agreement between the thresholds that an expert user would have applied and the software generated ones. The exceptions to this were caused by artifact formation in some wells. See supplementary Figure S.9 for an example. It should be noted that the QIAcuity runs was done over 4 day2s rather than 3 due to a technical interruption on day 3, i.e. instead of 2 runs on each day, day 3 had one run and day 4 had one run.

#### Optolane LOAA real-time dPCR

For a real-time dPCR system the LOAA system (Optolane Technologies Inc, Yongin-si, Gyeonggi-do, Republic of Korea) was used with the Dr. PCR cartridge. The final reaction volume was 30 µL. The LOAA system has 1 channel for excitation and 2 channels for scanning and relies on FRET probes for detection in the 2^nd^ scanning channel. Therefore, FRET FAM/Cy5 probes were designed for both the BPV-2 and IFN assays. Furthermore, instead of a single triplex reaction the assay was split into two – one with BPV 1 and 2 and another with IFN. The final reaction mix consisted of Dr. PCR mastermix 1X, BPV primers at 0.75 µM, the BPV-1 FAM probe at 0.2 µM, BPV-2 FRET probe at 0.8 µM. For the IFN reactions the primers were at 0.5 µM and the IFN FRET probe at 0.8 µM. 1.5 µL of DNA was loaded in both cases and the remaining volume was HPLC grade water. The cycling conditions were: 95 °C for 3 min, then 40 cycles of 95 °C for 15 s and 57 °C for 30 s, afterward an additional 5 cycles of 95 °C for 15 s and 55 °C for 30 s again using the touchdown approach described (24). Partition classification in the LOAA utilizes Cq values generated for each partition. After assessing different approaches, it was deemed that the platform software partition calling was the strongest option (Dr. PCR Analyzer 2), as such, the LOAA utilizes the built-in software for partition classification.

## Results and Discussion

### Cross-platform performance of the assays

#### Limit of blank (LOB)

Establishing the LOB is critical for characterizing an assay’s performance at low target concentrations. The LOB is the upper limit of false positive signals that can be observed in blank samples, i.e., samples not containing the analyte(s) of interest, effectively serving as the threshold that separates false positives from true positives (defined in Table 1 and the materials and methods section). Although in PCR, negative blanks should be aimed for, non-null blanks in dPCR may have various causes—assay design issues, cross-well contamination, instrument artifacts—which warrant careful evaluation (25). The full dilution series data for BPV-1 on the Naica platform (including 9–10 replicates per concentration level) is provided in Supplementary Table 2.S. This dataset illustrates the structure and granularity of the underlying data used for LOB, LOQ, and LOD calculations.

In this study, LOB values were derived from 20 dPCR or 60 qPCR measurements of blank samples. Table 2 summarizes the target-specific LOBs observed across instruments.

**Table 2.** Limit of blank (LOB) values for all platforms. Note that the high LOB observed for QIAcuity IFN is likely caused by artifacts in 1 well falsely inflating the LOB. Cp/rxn denotes the total number of DNA copies in the reaction.

| parameter | qPCR | Naica | QIAcuity | LOAA |
| --- | --- | --- | --- | --- |
| BPV-1 LOB | 0.42 cp/rxn | 0.00 cp/rxn | 0.00 cp/rxn | 2.45 cp/rxn |
| BPV-2 LOB | 0.00 cp/rxn | 1.02 cp/rxn | 0.72 cp/rxn | 0.89 cp/rxn |
| IFN LOB | 0.00 cp/rxn | 0.00 cp/rxn | 7.07 cp/rxn | 1.08 cp/rxn |

For BPV-1, Naica and Qiacuity each had an LOB of 0.00 copies/µL, in contrast to qPCR at 0.42 copies/µL and LOAA at 2.45 copies/µL. For BPV-2, qPCR’s LOB was 0.00 copies/µL, while Naica, QIAcuity, and LOAA had LOBs of 1.02, 0.72, and 0.89 copies/µL, respectively. With IFN, Naica and qPCR each had an LOB of 0.00 copies/µL, whereas QIAcuity and LOAA’s LOB rose to 7.07 copies/µL and 1.08 copies/µL, respectively. This illustrates how LOB values are both platform- and assay-dependent. While LOAA consistently showed non-zero LOBs across all targets, this suggests either residual background noise or an issue with the digital Cq classification approach used in the LOAA platform. Naica and QIAcuity only showed elevated LOBs for certain targets. Notably, IFN on QIAcuity had an unusually high LOB, which was caused by artifacts. Whereas such artifacts could be excluded through thresholding in most NTC wells, the Relative fluorescence units (RFU) values were too high to exclude in a single well.

The source of the false positive signal in dPCR reactions could either be caused by low levels of contamination with PCR product from previous reactions, off-target selectivity, or could be caused by a low level of artifacts in the dPCR partitions, causing false positive calls for some partitions (26). As contamination cannot be excluded, we advise running blank samples throughout the validation step, and not only at the start of the validation, as the risk of laboratory contamination increases with time. Accordingly, an LOB should be frequently confirmed.

#### Sensitivity and limit of detection (LOD)

Accurate determination of the LOD is crucial for comparing the analytical performance of PCR systems. In this study, two complementary approaches were used to estimate the LOD for each target (BPV-1, BPV-2, and IFN). The first approach, here referred to as the Type 1 (LOB-based) LOD, is the lowest concentration at which 95% of replicates are above the LOB. Since this method depends on measuring low-level dilution samples above the LOB, it can yield a more conservative—or higher—LOD when the LOB is non-zero or when the number of technical replicates is limited. As stated in ISO 20395:2019, the precision of the LOD estimate depends on both the number of replicates at each concentration and the spacing between dilutions. While the standard recommends at least 10 replicates per dilution, in practice this threshold requires all 10 replicates to be positive, effectively imposing a 100% positivity criterion (14). The CLSI EP17-A guideline provides a more practical framework, recommending at least 20 replicate measurements for verification and 60 for establishing LOB/LOD (22). Our design followed these recommendations, with 20 blanks for each dPCR platform and >60 blanks for qPCR, and >20 replicate measurements near the LOD. Although we did not explicitly analyze the effect of replicate number, it is important to note that using fewer replicates can yield unstable quantile or curve-fit estimates, particularly for empirical LOB and Type 2 LOD calculations. In contrast, the Type 2 LOD is derived from fitting a sigmoidal curve to the fraction of positives observed at all dilution levels, after which the concentration at which 95% of replicates are expected to be positive is estimated. By considering the entire dataset, the Type 2 approach often provides a lower and potentially more precise estimate. However, LOD estimates must remain physically plausible: Poisson statistics indicate that achieving ≥95% detection requires an average of ∼3 target copies per reaction, setting a theoretical lower bound. In our dataset, the Qiacuity IFN assay illustrates this limitation — the Type 2 LOD (1.47 copies per reaction (cp/rxn)) was lower than both the measured LOB (7.07 cp/rxn) and the Poisson limit (table 3). This may occur because the curve-fit method does not explicitly account for non-null blank measurements and can therefore return unrealistically low values. Another methodological consideration is that we maintained a constant input volume of 1.5 µL across all platforms. In principle, platform sensitivity would be best evaluated by maximizing the sample input permitted by each system, as larger reaction volumes generally allow for lower theoretical LODs. However, the usable input volume is assay dependent (e.g., influenced by mastermix formulation and multiplexing requirements), and in clinical practice sample material is often limited, making such maximization impractical. By fixing the input at 1.5 µL—the volume routinely used for this assay in its established qPCR implementation—we benchmarked the platforms under conditions aligned with actual diagnostic practice for this specific assay, at the expense of not assessing their maximal theoretical sensitivity.

**Table 3.** The LOD Type 1 and 2 values for all assays and PCR platforms. Cp/rxn denotes the total number of DNA copies in the reaction.

| LOD Type 1 | BPV-1 | BPV-2 | IFN |
| --- | --- | --- | --- |
| **qPCR** | 49.265 cp/rxn | 11.861 cp/rxn | 9.9 cp/rxn |
| **Naica** | 29.1561 cp/rxn | 33.167 cp/rxn | 21.48 cp/rxn |
| **QIAcuity** | 6.47 cp/rxn | 35.64 cp/rxn | 45.92 cp/rxn |
| **LOAA** | 6.47 cp/rxn | 7.13 cp/rxn | 22.96 cp/rxn |

| LOD Type 2 | BPV-1 | BPV-2 | IFN |
| --- | --- | --- | --- |
| qPCR | 15.85 cp/rxn | 3.25 cp/rxn | 3.46 cp/rxn |
| Naica | 3.04 cp/rxn | 5.14 cp/rxn | 6.53 cp/rxn |
| QIAcuity | 3.06 cp/rxn | 6.38 cp/rxn | 1.47 cp/rxn |
| LOAA | 4.15 cp/rxn | 4.81 cp/rxn | 7.41 cp/rxn |

Table 3 presents the Type 1 and 2 LOD values for each platform, while Figure 1 illustrates the Type 2 LOD estimates. In general, Type 1 LOD estimates are higher across all platforms and targets. For BPV-1, the difference between Type 1 and Type 2 is particularly striking for the Naica and QIAcuity dPCR systems, which show the Type 1 LODs ranging from 6.47 to 29.16 copies per reaction, but much lower Type 2 LODs in the range of 3.04 to 4.03 copies per reaction. This pattern underscores the more conservative nature of the LOB-based approach. A similar trend appears for qPCR with BPV-1, where the Type 1 LOD (49.27 copies per reaction) is substantially higher than the Type 2 LOD (16.42 copies per reaction). Although the Type 1 LOD ensures that the assay can truly distinguish signal from blank at the chosen concentration, it can depend on how that concentration is selected and on the number of replicates available.

**Figure 1.**
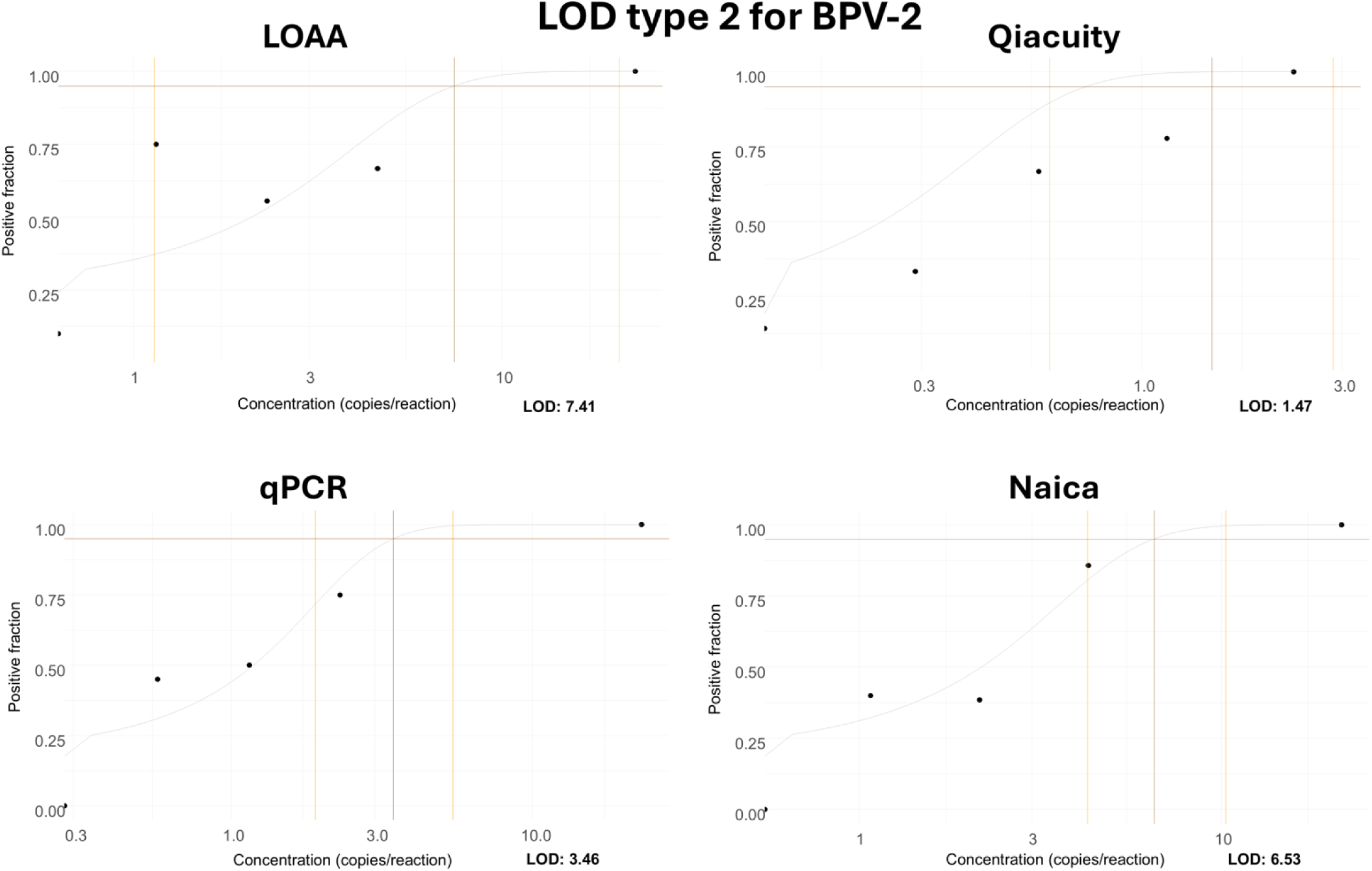
example of type 2 LOD data output from PCR-ValiPal. A logistic model is fit to the observed fractions of positives (grey curve) using the full set of replicates (n≈9 per dilution point), the latter obtained by calculating the number of positives divided by the number of observations at each dilution level. The type 2 LOD is then obtained as the crossing point of that logistic curve with the user-provided LOD threshold. A confidence interval for the LOD is obtained by a parametric bootstrap procedure (yellow horizontal lines). Briefly, for each dilution point, the fraction of positives is obtained by sampling from a binomial distribution with parameters p and n, where p is the observed fraction of positives for that dilution level, and n the total number of observations for that dilution level. A logistic curve is fit to each bootstrapped dilution curve, and a 95% percentile confidence interval calculated. The vertical brown lines indicate the 95% positive fraction, the type 1 LOD is the first dilution step above this line.

When looking at BPV-2, the Type 1 LOD for QIAcuity (35.64 copies per reaction) sits well above its Type 2 counterpart (6.38 copies per reaction). In contrast, qPCR shows a relatively modest gap between Type 1 and Type 2 (11.86 vs. 5.42 copies per reaction). The same pattern emerges for IFN, most notably with QIAcuity, which has a Type 1 LOD of 45.92 copies per reaction but a much lower Type 2 LOD of 2.75 copies per reaction. These discrepancies highlight that when an assay has a non-zero LOB or frequently yields positives in blank samples, a sigmoidal-fit approach that does not explicitly account for background noise can produce misleading LOD estimates—potentially inflating or deflating the threshold, depending on the dilution-specific fraction of positives. Conversely, although having a negligible LOB often allows the Type 1 and Type 2 methods to converge, this is not guaranteed, as the steepness and variability of the detection curve can still drive large differences (e.g., QIAcuity for BPV-1). Hence, LOB is only one among several factors—alongside replicate count, dilution spacing, and the assay’s signal kinetics—that influence the degree of agreement between model-based and empirical LOD estimates. Overall, these results illustrate the strengths and limitations of each approach. The LOB-based method is straightforward when the blank is not zero, but in practice, it often overestimates the LOD if the dilutions tested sit at a level where variability or replicate count can affect the outcome, or if dilutions are relatively widely spaced. The sigmoidal-curve-fit method makes better use of the complete dilution series, so its estimate will usually be more precise in large datasets, yet it may produce an unrealistically optimistic LOD if blank measurements are non-zero.

In this study, the Type 1 LOD was determined using between 9 and 12 replicates per dilution level, with the exact number varying by concentration and platform, typically highest near the expected LOD.(14). Under these conditions, the required 95% positivity threshold translates to detecting the target in all 10 replicates (100% positive). This means that, in practice, the Type 1 LOD is based on complete detection rather than allowing for a small fraction of false negatives. Researchers aiming to define the Type 1 LOD using a 5% α threshold (i.e., allowing 95% positivity with margin for false negatives) would require at least 20 replicates per concentration level to achieve the necessary resolution.

Taken together, these findings underscore the importance of interpreting LOD estimates in the context of actual blank measurements, replicate numbers, and assay characteristics. Relying solely on the Type 1 or Type 2 definition can be misleading. Systems that produce a measurable blank signal (whether due to contamination, assay design, or instrument artifacts) may favor the Type 1 approach to avoid undue optimism at low concentrations. Conversely, laboratories with extensive dilution series data and truly zero-signal blanks may find the Type 2 LOD to be a good reflection of practical assay sensitivity. In most cases, examining both LOD definitions helps researchers appreciate the range of likely detection thresholds and choose the more suitable metric for their application.

#### Limit of quantification (LOQ)

In contrast to the LOD, which serves as a threshold to detect the presence of analyte, the LOQ specifies the lowest concentration at which measurements can be quantified with acceptable precision (14). It is important to note that the required level of precision is not universally fixed. Rather, it should be defined by the user based on practical, regulatory, or clinical considerations. In this study, LOQ values were determined by testing reference concentrations and evaluating variation, using a coefficient of variation (CV) of 30% as the cutoff value. The LOQ data appears in Table 4.

**Table 4.** The LOQ values for all platforms. Cp/rxn denotes the total number of DNA copies in the reaction.

| parameter | qPCR | Naica | QIAcuity | LOAA |
| --- | --- | --- | --- | --- |
| BPV-1 LOQ | 985.31 cp/rxn | 29.16 cp/rxn | 32.36 cp/rxn | 647.00 cp/rxn |
| BPV-2 LOQ | 1186.12 cp/rxn | 33.32 cp/rxn | 35.54 cp/rxn | 713.00 cp/rxn |
| IFN LOQ | 99.00 cp/rxn | 42.97 cp/rxn | 918.58 cp/rxn | 459.29 cp/rxn |

Because samples with low concentrations are typically associated with decreased measurement precision, the LOQ is defined as the lowest concentration level at which a sufficiently high (relative) precision is obtained. This is usually determined by a maximum acceptable CV, such as 30%. Samples below this threshold may still be detected (above the LOD) but cannot be reliably quantified, i.e. the variance is higher than the chosen threshold. Moreover, although dPCR inherently offers absolute quantification without relying on external standards, laboratories often adopt user-defined CV cutoffs to satisfy clinical or regulatory constraints. These cutoffs help account for subtle technical variances—such as pipetting variation, partition uniformity, droplet stability—that even an “absolute” method like dPCR does not automatically eliminate. It is also worth noting that recommended CV thresholds can vary among research and diagnostic fields, highlighting the importance of empirically validating any chosen cutoff to ensure it aligns with each specific assay’s requirements.

For BPV-1, the qPCR LOQ (985.31 copies/reaction) was substantially higher than Naica (29.16 copies/µL) and QIAcuity (32.36 copies/reaction), suggesting that qPCR is less reliable at lower concentrations for this target. For BPV-2, qPCR again showed the highest LOQ (1186.12 copies/µL), whereas Naica and QIAcuity measured in the 30–36 copies/reaction range. With IFN, Naica displayed the lowest LOQ (42.97 copies/reaction), followed by qPCR at 99 copies/reaction, LOAA at 459.29 copies/reaction, and QIAcuity at 918.58 copies/reaction—indicating that QIAcuity requires higher IFN concentrations for reliable quantification. It should be noted that for the QIAcuity platforms, one dilution series had significantly lower concentration than the replicates for the other days (e.g. average for dilution step 7 day 1-3 = 94.52, average day 4 = 57.76). There is no experimental basis for excluding these replicates as encompassing all variation across dilution series was an aim for all platforms. However, without this dilution series, the LOQ for IFN QIAcuity becomes 91.86 copies/reaction (the next dilution step), comparable to the qPCR. This also highlights that the LOQ estimation has similar vulnerabilities as the type 1 LOD. For instance, in our data, the dilution series has a 1:10 dilution step, from 918.58 copies/rxn to 91.858 copies/rxn. The CV for the 918.58 copier/rxn was 33%, putting it just above the LOQ threshold. This means that the current approach to LOQ determination gives a conservative estimate.

#### Linearity

Linearity was assessed for each assay primarily through quadratic regression, with additional confirmation via r² values derived from robust weighted least squares (WLS). Following the approach recommended in dPCalibRate (15), the quadratic term provides a sensitive and specific test for deviations from linearity, while r² serves as a useful but less critical supporting measure. This combined strategy allows a more rigorous delineation of the assay’s effective dynamic range.

The quadratic regression evaluates whether introducing a second order (curvature) term significantly improves the model fit. A significant quadratic term suggests the presence of curvature—either upward or downward—across the concentration range, thus signaling a deviation from strict proportionality.

qPCR exhibited significant non-linearity for BPV-1 (*p = 3.0e-2*), confirming that concentration-response behavior deviated from ideal linearity for this target (Table 5). Although the r² for BPV-1 on qPCR (0.890) also indicated a poor fit, the primary evidence of non-linearity comes from the significant quadratic term. Naica likewise displayed borderline non-linearity for BPV-1 (*p = 4.0e-2*) despite a high r² (0.983), underscoring that a strong r² alone can mask subtle but statistically significant curvature.

**Table 5.** Results of statistical linearity tests. Overall, the quadratic regression analysis highlighted specific instances of non-linearity that would have been understated or missed by r² assessment alone. In particular, qPCR for BPV-1 and Naica for BPV-2 exhibited significant curvature despite generally acceptable r² values. This pattern emphasizes the critical role of formal statistical testing for curvature when evaluating assay performance, especially when working across broad dynamic ranges or platforms susceptible to subtle amplification biases. Where both the quadratic term and r² agreed (as seen with IFN assays), confidence in the assay’s linearity was correspondingly high.

| target | metric | qPCR | Naica | QIAcuity | LOAA |
| --- | --- | --- | --- | --- | --- |
| **BPV-1** | quadratic p | 3.2e-2 | 4.2e-2 | 6.1e-1 | 6.8e-2 |
| | $r^2$ (robust WLS) | 0.889 | 0.983 | 0.980 | 0.968 |
| **BPV-2** | quadratic p | 3.8e-1 | 3.8e-2 | 3.8e-1 | 5.0e-1 |
| | $r^2$ (robust WLS) | 0.935 | 0.984 | 0.986 | 0.966 |
| **IFN** | quadratic p | 6.6e-1 | 6.0e-2 | 4.6e-1 | 1.4e-1 |
| | $r^2$ (robust WLS) | 0.974 | 0.989 | 0.978 | 0.983 |

For BPV-2, most platforms maintained non-significant quadratic terms (*p > 5.0e-2*), except for Naica, where *p = 4.0e-2* suggested a mild but statistically significant deviation from linearity. This finding, again, was not readily apparent from the corresponding high r² values, which remained above 0.96 across all systems.

IFN consistently demonstrated the most robust linear behavior across platforms, with relatively high *p*-values for the quadratic term (all *p > 6.0e-2*) and relatively high r² values (>0.97). These results indicate that IFN measurements preserved proportionality throughout the concentration range, with no detectable curvature effects.

#### Deviation from fit

To complement the statistical tests, PCR-ValiPal calculates the percentage deviation from the linear model for each concentration level (Figure 2). Negative values signify underestimation, while positive values indicate overestimation relative to the best-fit line.

**Figure 2.**
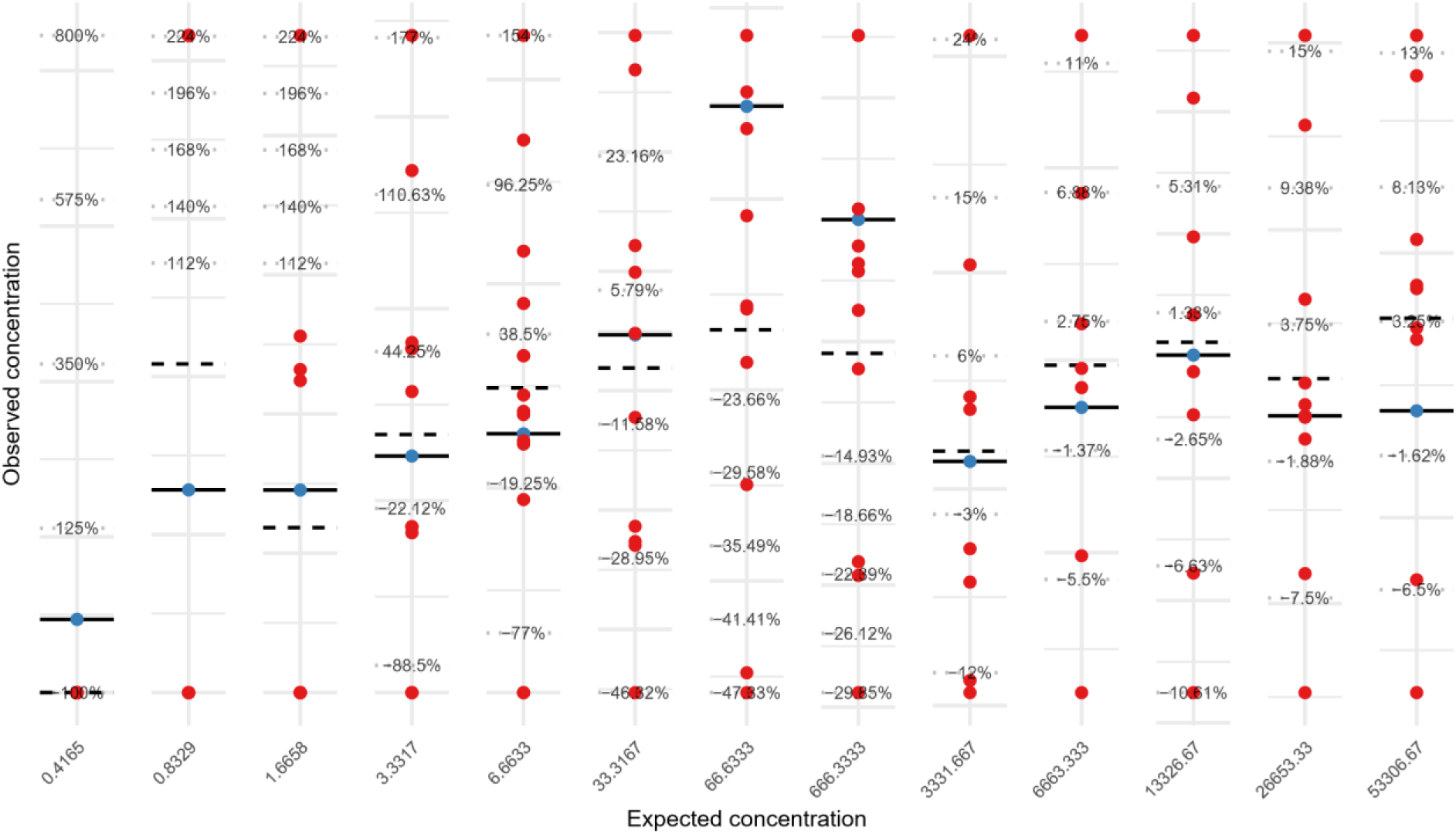
Example of deviation form fit output from PCR-ValiPal. Such a figure has been generated for all platform/assay combinations. Observed (red dots, n≈9 per dilution point), average of observed (dashed line) and fitted (blue dots, full line) values. For each dilution level, the deviation from the expected (fitted) value is visualized. Large deviations between (average) of observed values and the fitted value indicate deviation from linearity. This figure was generated for each platform/assay combination, the above is an example of the data.

This data highlighted larger deviations for qPCR in some BPV-1 and BPV-2 measurements, aligning with the lower r² values for those targets. By contrast, Naica and QIAcuity generally remained within ±15% of the fitted curve—except for certain low-concentration points—consistent with their higher r² scores. LOAA typically displayed moderate deviations (±20%) but remained fairly stable across mid-to-high concentrations. The spread of residuals increased at lower concentrations, consistent with the presence of heteroscedasticity in PCR (Figure 2).

#### Trueness analysis

Trueness is the deviation of observed results from expected values, sometimes called bias (14). It can reveal systematic errors, such as under- or overestimation due to dilution inaccuracies, instrument bias, or suboptimal assay design. High levels of precision do not necessarily guarantee high trueness—an assay can be consistently ‘off’ from the true value.

Trueness was evaluated and compared across systems (see Figure 3 for example data). Negative values indicate underestimation, while positive values denote overestimation of the expected concentration.

**Figure 3.**
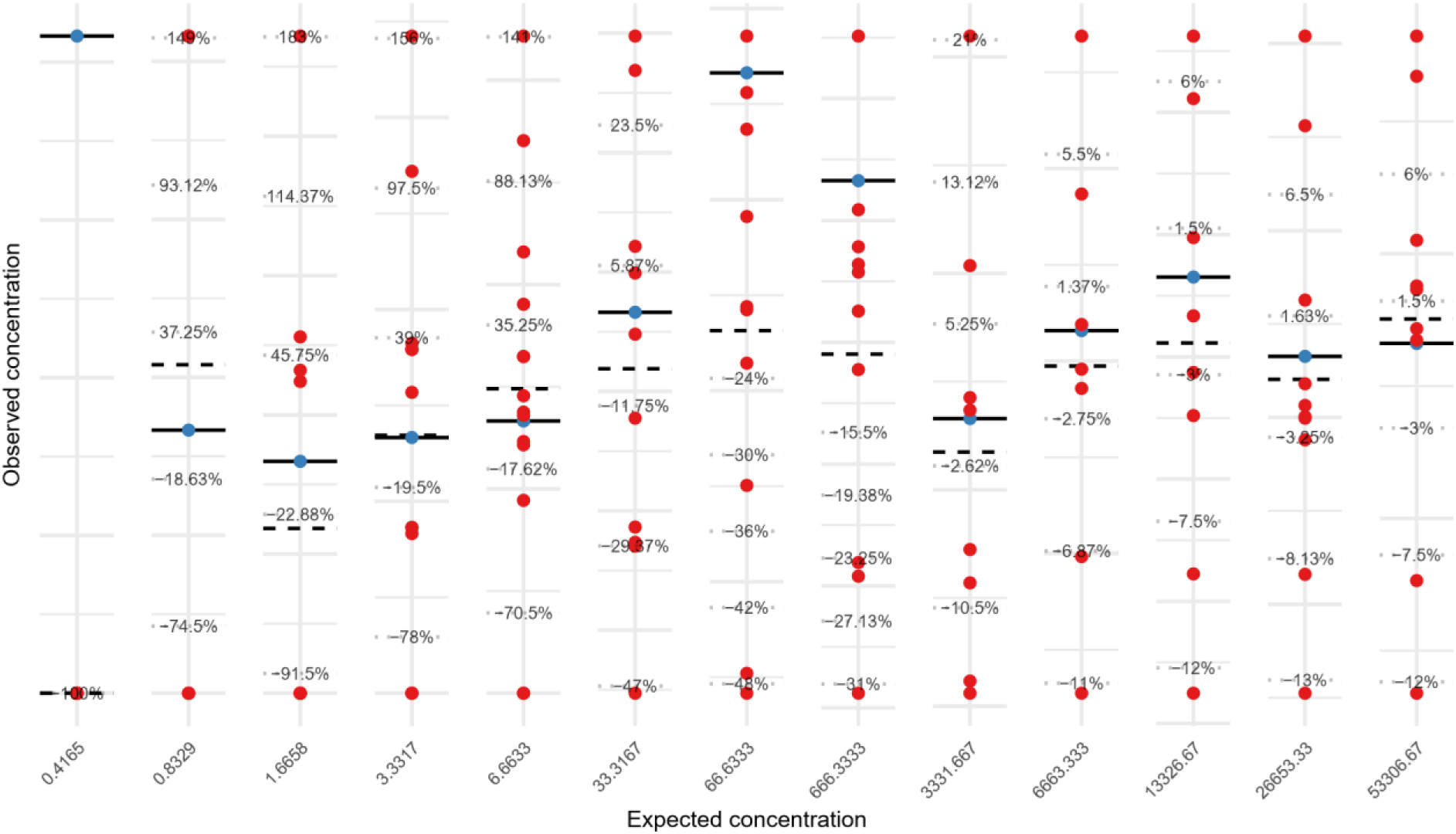
Observed (red dots), average of observed (dashed line) and expected (blue dots, full line) values. For each dilution level, the deviation from the expected (fitted) value is visualized. Large deviations between (average) of observed values and the fitted value indicate deviation from linearity. This figure was generated for each platform/assay combination, the above is an example of the data.

These values illustrate that Naica and QIAcuity often hovered near single-digit deviations from the expected concentration, while LOAA demonstrated a consistent negative bias. qPCR showed somewhat larger swings for the BPV targets, although it performed acceptably for IFN, staying mostly within ±10% at mid- to high-level concentrations.

In particular, the consistent negative bias displayed by LOAA suggests that, while the system underestimates true concentrations, it does so in a reproducible manner. This predictable offset can often be compensated by applying a calibration factor or correction curve, provided that the bias remains stable across multiple runs and concentrations. While these deviations largely reflect inherent assay and platform behaviors, a final methodological limitation also warrants consideration.

We note that the standard curve for qPCR was generated using Naica-derived concentrations. Because the absolute quantification used for qPCR calibration was itself obtained on the Naica platform, this introduces a degree of circular reasoning when comparing qPCR to dPCR performance. While Naica’s digital PCR quantification is assumed to be accurate, any systematic bias in its measurement — whether due to partition classification, droplet volume estimation, or assay behavior — would propagate to the qPCR standard curve and thus potentially mask differences between platforms. This limitation should be considered when interpreting the relative agreement between qPCR and dPCR results. Ideally, an orthogonal method would be used to generate the reference material, decoupling the calibration process from any individual system.

A closer examination of the LOAA results reveals that while resolution (the ability to distinguish positive and negative partitions) are comparable to those of QIAcuity, misclassification of partitions based on amplification curves contributes to the observed negative bias. Specifically, a portion of partitions falling within the positive fluorescence cloud are incorrectly classified as negative, likely due to the system’s reliance on Cq-based thresholding. This issue is most evident in partitions with low Cq values but strong endpoint signals—patterns typically called positive in dPCR platforms but missed by Cq-based classification. Although threshold adjustments are possible, overly permissive settings risk introducing false positives in the negative cluster.

A final note concerns the commonly implemented yet discouraged strategies (e.g., relying solely on simple r² or unweighted regressions to evaluate linearity). These methods are included in the PCR-ValiPal output, not to endorse their use, but rather to provide a point of comparison against more robust approaches. By making both traditional and recommended strategies accessible, users can directly observe how different analytical choices influence parameter estimates, facilitating a smoother transition toward standardized best practices.

## Conclusion

Taken together, these findings demonstrate that overall performance varies across platforms and is highly dependent on the specific target assay. Both dPCR systems (QIAcuity and Naica) achieve strong sensitivity (particularly for BPV-1 and IFN), low LOB and LOQ values, and excellent linearity, all while displaying small deviations from expected concentrations—indicating a high degree of accuracy. LOAA likewise exhibits strong linearity and consistently excels in within-run precision; however, it tends to underestimate target concentrations by a predictable margin. qPCR proves most sensitive for BPV-2 but is otherwise more variable, showing relatively larger deviations and inconsistent detection at very low concentrations, especially for BPV-1. Nonetheless, qPCR remains sufficiently accurate for IFN and demonstrates reasonable performance for BPV-2.

In sum, while dPCR and real-time dPCR generally maintain tighter precision, lower variability, and smaller systematic biases, qPCR can still be effective if properly calibrated and validated for each analyte. These results underscore the value of method-specific optimization and highlight that “best” performance cannot be assumed across all targets without empirical verification. Notably, although discouraged strategies remain part of the PCR-ValiPal output for comparative purposes, the recommended methods should be prioritized to ensure accurate and reproducible assay validation. For this purpose, the PCR-ValiPal Shiny application is freely available at: https://digpcr.shinyapps.io/valipal/

## Supplementary information

### Oligonucleotides

#### S.1 gBlock sequences

##### BPV1

atgaatcgggtgagcaaccttttaatattactgatgcagattggaaatctttttttgtaaggttatgggggcgtttagacctgattgacgaggaggaggatagtgaagaggatggagacagcatgc

##### BPV2

atgaatcgggtgagcaaccttttactattactgatgcagattggaaatctttttttgtaaggttatgggggcgcttagacctggttgacgaggaggaggatagtgaagaggatggagacagcatgc

##### IFN fragment

Ctgttcctgtcttcatcatgacctacaggtggatcctcccaatggccctcctgctgtgtttctccaccacggctctttctgtgaactatgacttgcttcggtcccaactaagaagcagcaattcag

#### Primers and probes

**Table S.1.**
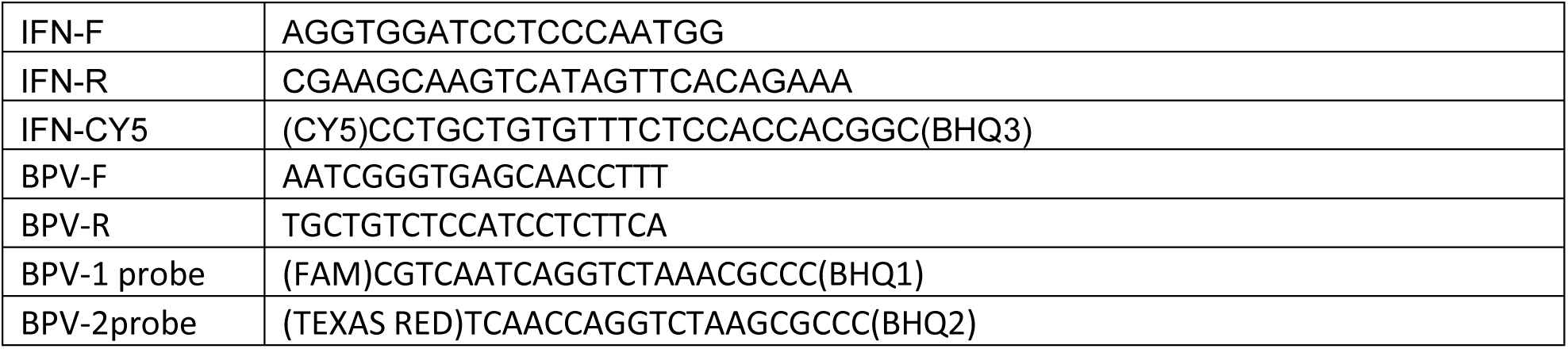
Primer and probe sequences for BPV-1 and 2. Note that for Optolane FRET FAM/CY5 are used.

### One-color plot examples

The following are 1-color plots from the various platforms for all assays.

BPV-1 is always FAM, BPV-2 is HEX/VIC in all platforms except LOAA where it is FRET-Cy5 and IFN is always Cy5

**Figure S.1.**
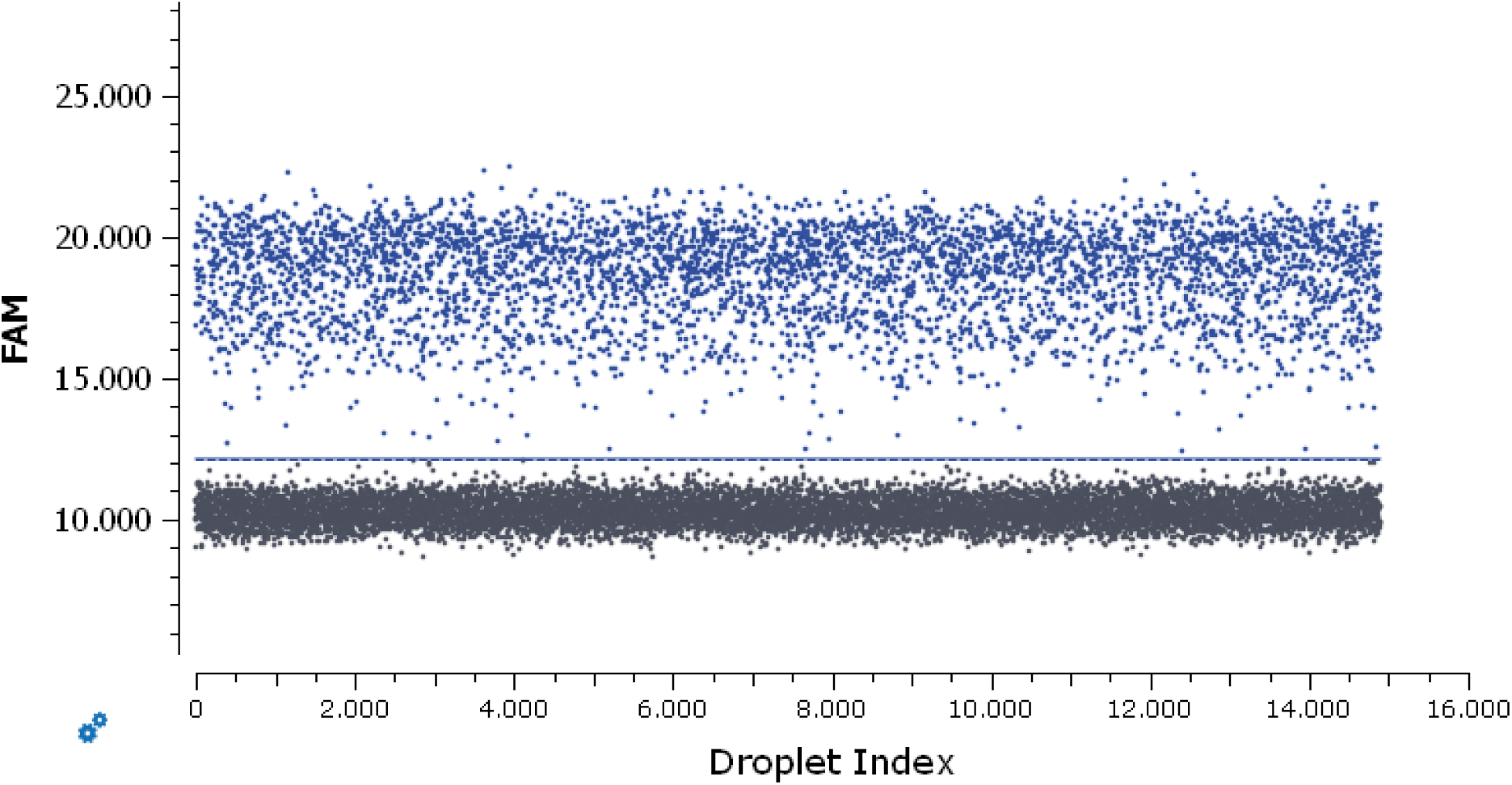
Example of BPV-1 1-color plot from the Naica platform.

**Figure S.2.**
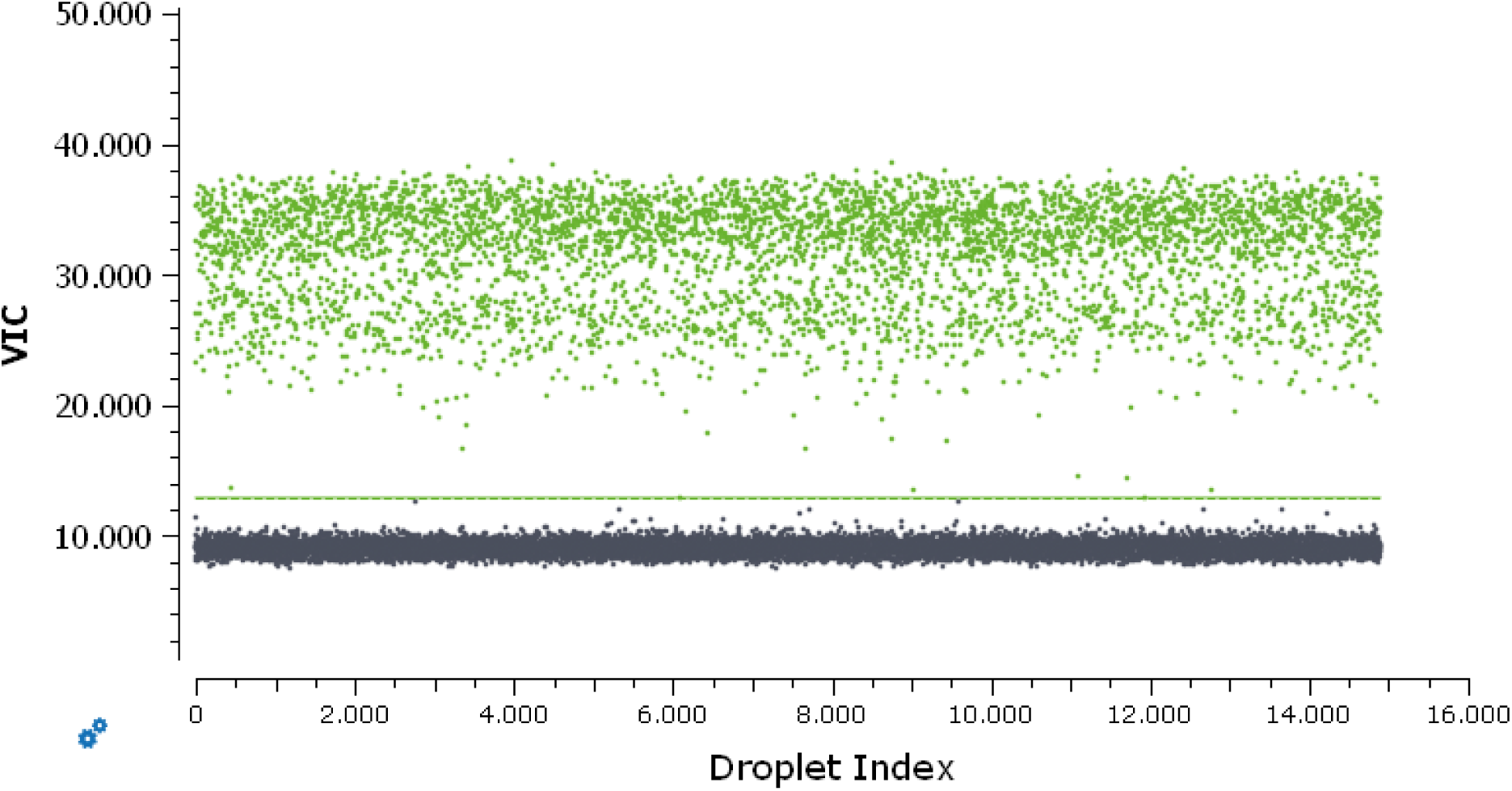
Example of BPV-2 1-color plot from the Naica platform.

**Figure S.3.**
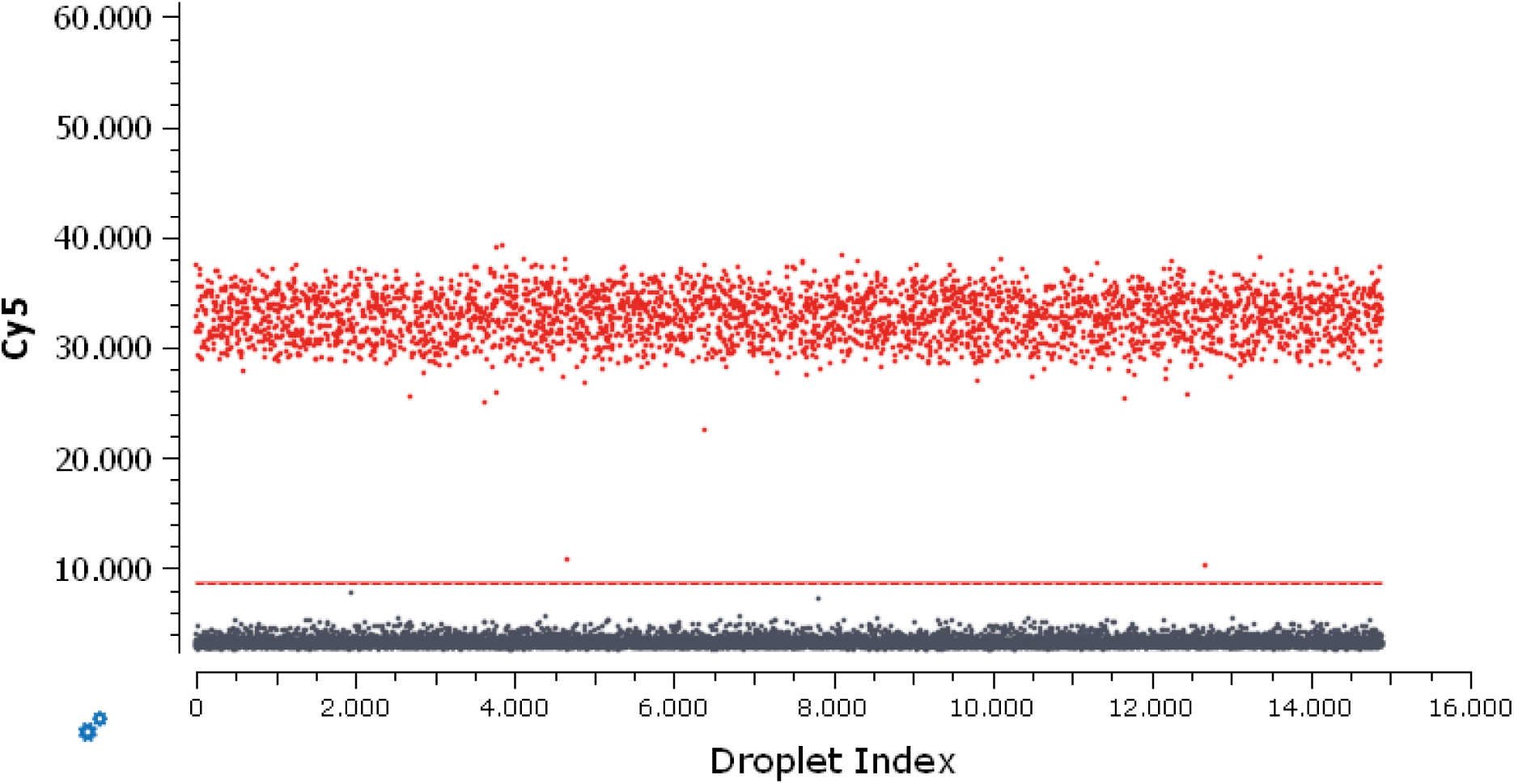
Example of IFN 1-color plot from the Naica platform.

**Figure S.4.**
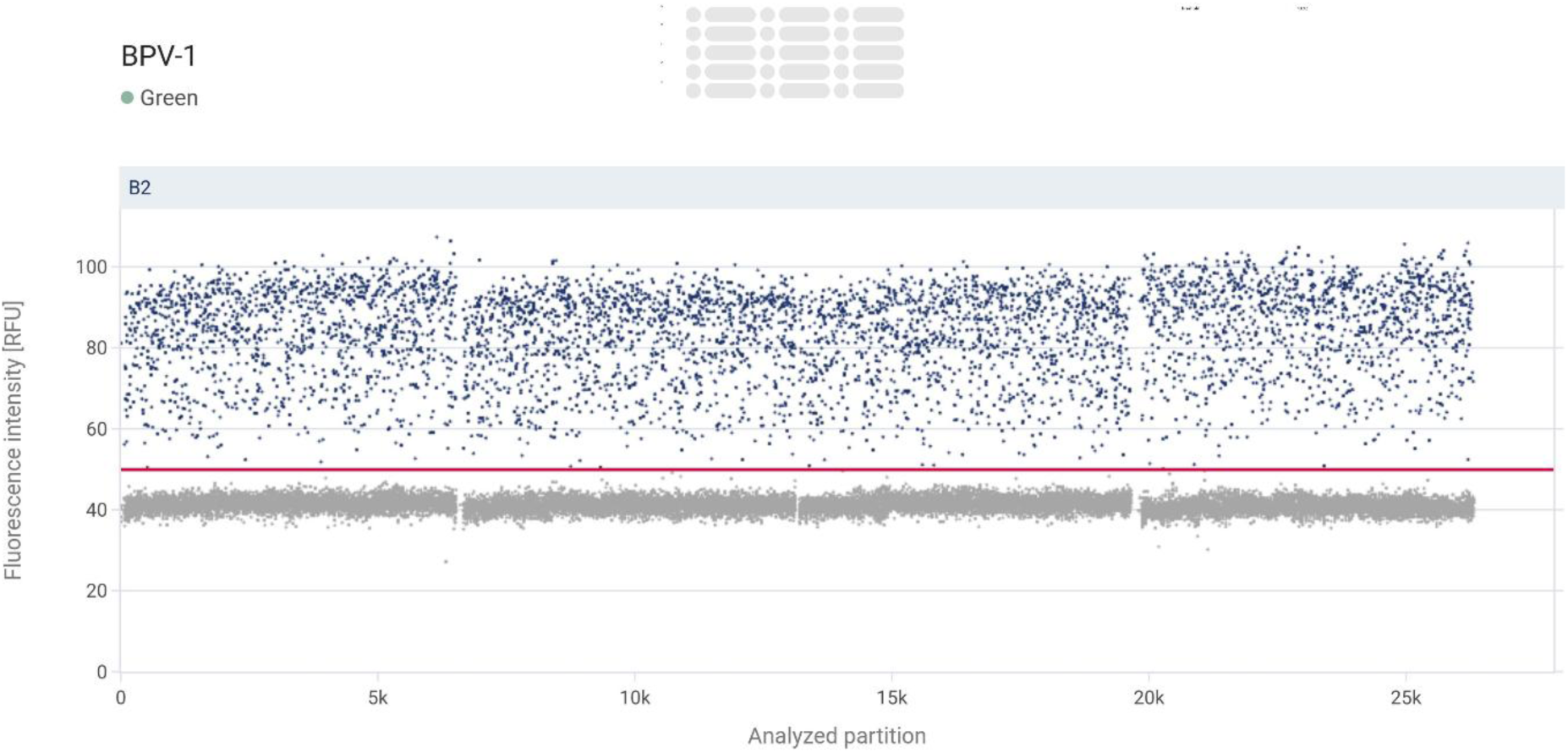
Example of BPV-1 1-color plot from the QIAcuity platform.

**Figure S.5.**
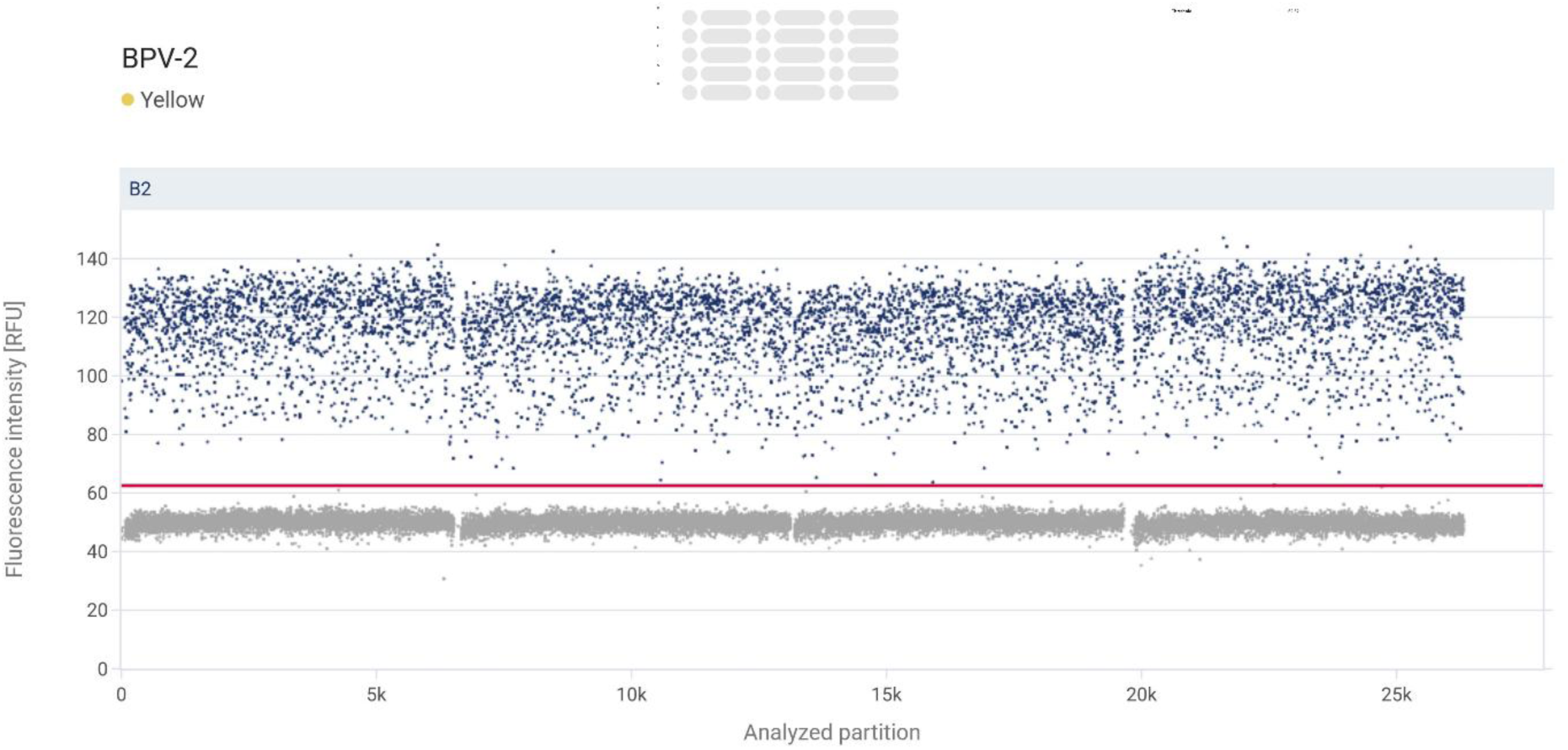
Example of BPV-2 1-color plot from the QIAcuity platform.

**Figure S.6.**
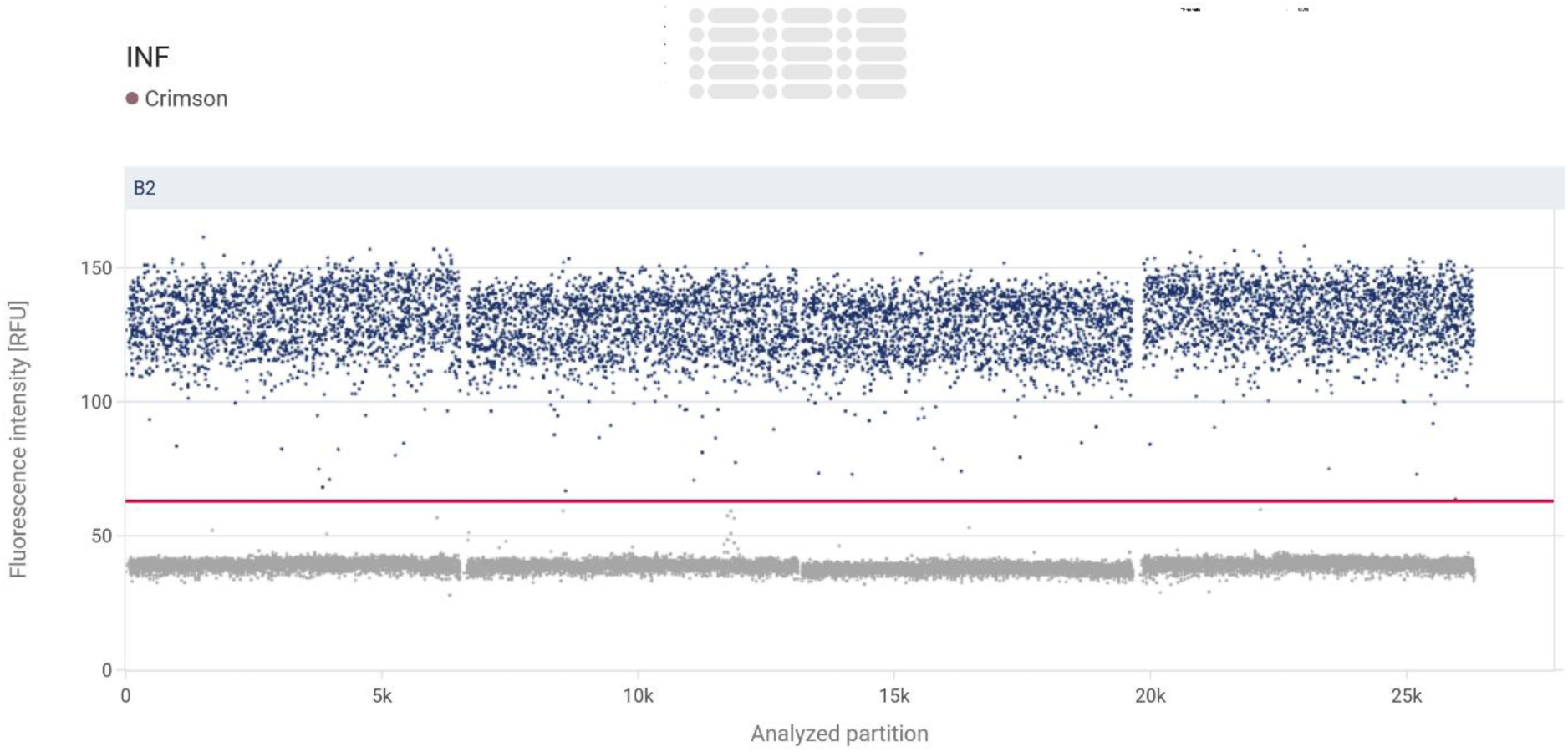
Example of IFN 1-color plot from the QIAcuity platform.

**Figure S.7.**
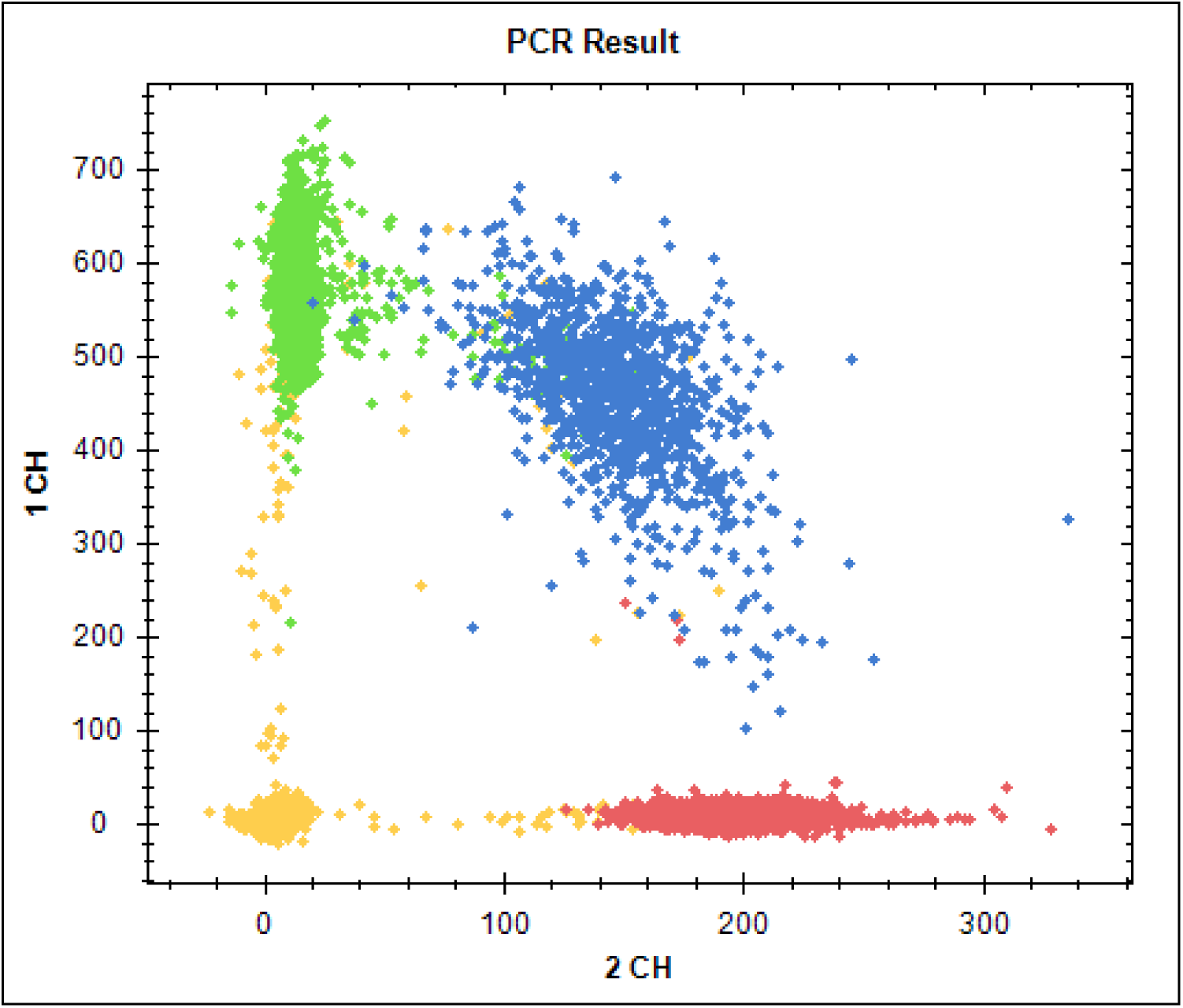
Example of BPV-1 and BPV-2 1-color plot from the LOAA platform.

**Figure S.8.**
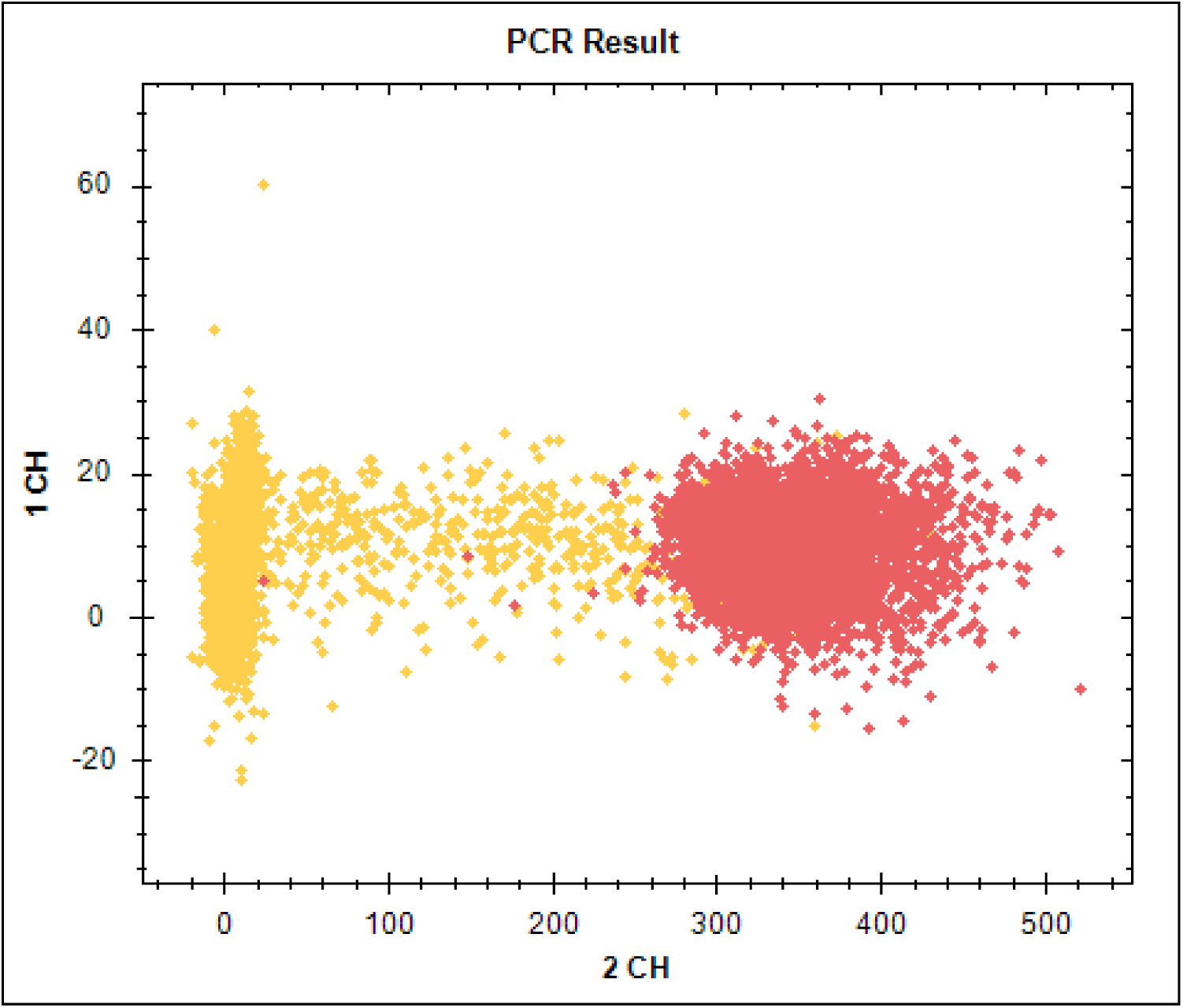
Example of IFN 1-color plot from the LOAA platform.

### Expected values for all assays

**Table S.2.** expected values for all assays and platforms.

| Naica | BPV-1 | BPV-2 | IFN |
| --- | --- | --- | --- |
| c1 | 0,364451 | 0,416458 | 0,268549 |
| c2 | 0,728903 | 0,832917 | 0,537097 |
| c3 | 1,457806 | 1,665833 | 1,074194 |
| c4 | 2,915611 | 3,331667 | 2,148389 |
| c5 | 5,831222 | 6,663333 | 4,296778 |
| c6 | 29,15611 | 33,31667 | 21,48389 |
| c7 | 58,31222 | 66,63333 | 42,96778 |
| c9 | 583,1222 | 666,3333 | 429,6778 |
| c10 | 2915,611 | 3331,667 | 2148,389 |
| c11 | 5831,222 | 6663,333 | 4296,778 |
| c12 | 11662,44 | 13326,67 | 8593,556 |
| c13 | 23324,89 | 26653,33 | 17187,11 |
| c14 | 46649,78 | 53306,67 | 34374,22 |
| qPCR | BPV-1 | BPV-2 | IFN |
| C1 | 0,615822 | 0,741323 | 0,618779 |
| C2 | 1,231643 | 1,482646 | 1,237559 |
| C3 | 2,463287 | 2,965293 | 2,475118 |
| C4 | 4,926574 | 5,930585 | 4,950235 |
| C5 | 9,853148 | 11,86117 | 9,90047 |
| C6 | 49,26574 | 59,30585 | 49,50235 |
| C7 | 98,53148 | 118,6117 | 99,0047 |
| C8 | 492,6574 | 593,0585 | 495,0235 |
| C9 | 985,3148 | 1186,117 | 990,047 |
| C10 | 4926,574 | 5930,585 | 4950,235 |
| C11 | 9853,148 | 11861,17 | 9900,47 |
| C12 | 19706,3 | 23722,34 | 19800,94 |
| C13 | 39412,59 | 47444,68 | 39601,88 |
| C14 | 78825,18 | 94889,37 | 79203,76 |
| C15 | 246328,7 | 296529,3 | 247511,8 |
| C16 | 492657,4 | 593058,5 | 495023,5 |
| C17 | 985314,8 | 1186117 | 990047 |
| Qiacuity | BPV-1 | BPV-2 | IFN |
| c1 | 0,4 | 0,45 | 0,58 |
| c2 | 0,81 | 0,89 | 1,14 |
| c3 | 1,62 | 1,78 | 2,3 |
| c4 | 3,24 | 3,56 | 4,6 |
| c5 | 6,47 | 7,13 | 9,18 |
| c6 | 32,36 | 35,64 | 45,92 |
| c7 | 64,72 | 71,28 | 91,86 |
| c9 | 647,15 | 712,78 | 918,58 |
| c10 | 3235,75 | 3563,88 | 4592,88 |
| c11 | 6471,5 | 7127,75 | 9185,76 |
| c12 | 12943 | 14255,5 | 18371,5 |
| c13 | 25886 | 28511 | 36743 |
| c14 | 51772 | 57022 | 73486 |
| LOAA | BPV-1 | BPV-2 | IFN |
| c1 | 0,4 | 0.45 | 0.29 |
| c2 | 0,81 | 0.89 | 0.57 |
| c3 | 1,62 | 1.78 | 1.15 |
| c4 | 3,24 | 3.56 | 2.30 |
| c5 | 6,47 | 7.13 | 4.59 |
| c6 | 32,36 | 35.64 | 22.96 |
| c7 | 64,72 | 71.28 | 45.93 |
| c9 | 647,15 | 712.78 | 459.29 |
| c10 | 3235,75 | 3563.88 | 2296.44 |
| c11 | 6471,5 | 7127.75 | 4592.88 |
| c12 | 12943 | 14255.50 | 9185.75 |
| c13 | 25886 | 28511.00 | 18371.50 |
| c14 | 51772 | 57022.00 | 36743.00 |

**Table S.3.** Overview of dilution series points used for assay validation.

|  |  |
| --- | --- |
| C17.5 | Platform stock |
| **C17** | 1/1,2 dilution of C17.5: |
| **c16** | 1/2 dilution of C17: |
| **c15** | 1/2 dilution of C16: |
| **c14** | 1/3,125 dilution of C15: |
| **c13** | 1/2 dilution of C14: |
| **c12** | 1/2 dilution of C13: |
| **c11** | 1/2 dilution of C12: |
| **c10** | 1/2 dilution of C11: |
| **c9** | 1/5 dilution of C10: |
| **c8** | 1/2 dilution of C9: |
| **c7** | 1/5 dilution of C8: |
| **c6** | 1/2 dilution of C7: |
| **c5** | 1/5 dilution of C6: |
| **c4** | 1/2 dilution of C5: |
| **c3** | 1/2 dilution of C4: |
| **c2** | 1/2 dilution of C3: |
| c1 | 1/2 dilution of C2: |

**Figure S.9.**
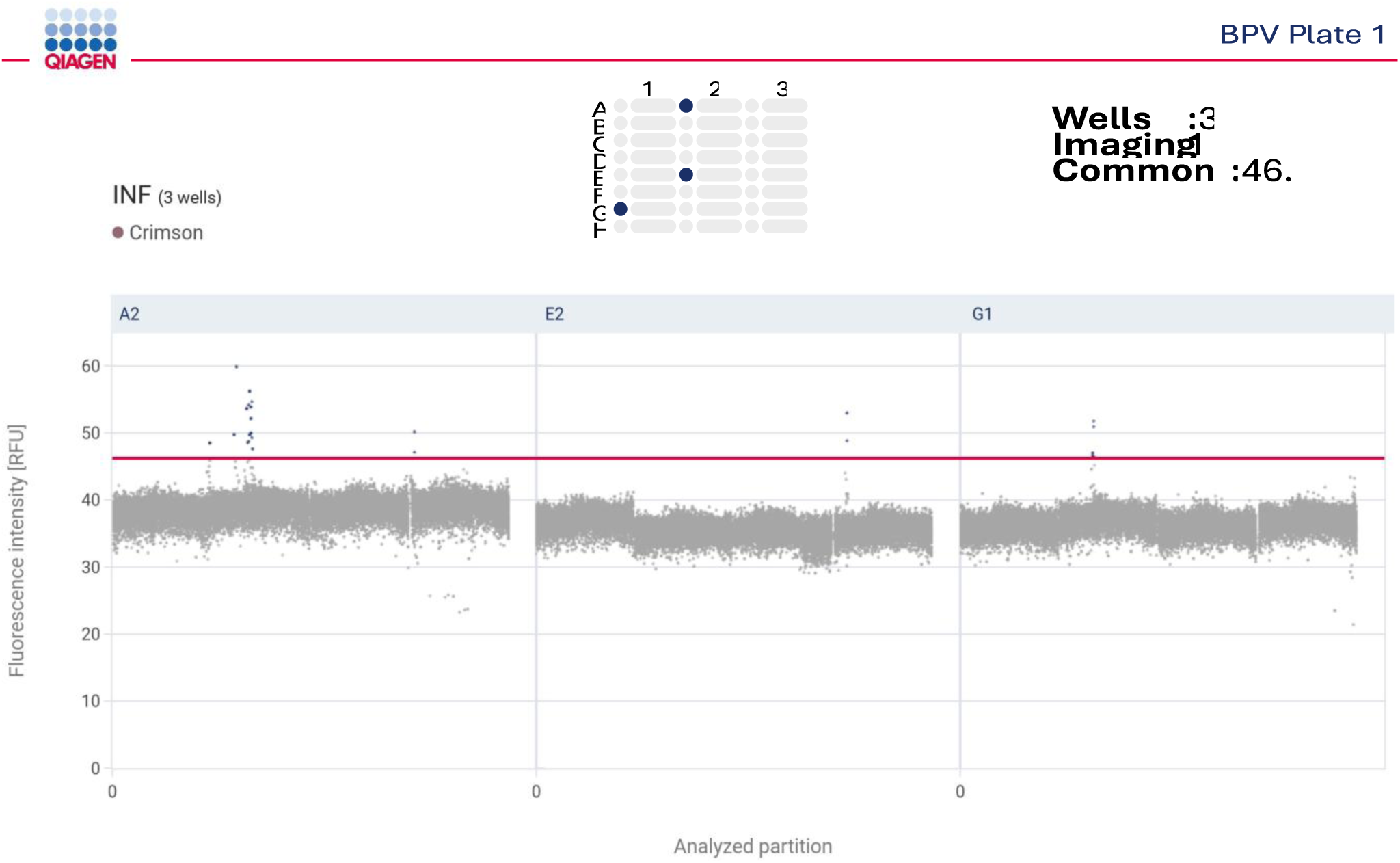
example of artifacts in Qiacuity NTC’s. these were only observed in the Cy5 channel.

**Table S.4.** Mean lambda (λ) values ± standard deviation (SD) for BPV-1, BPV-2, and IFN assays measured across three dPCR platforms (Naica, QIAcuity, LOAA) at each dilution level. Values are reported as λ (mean ± SD).

BPV-1
| Dilution | Naica ($\lambda \pm$ SD) | QIacuity ($\lambda \pm$ SD) | LOAA ($\lambda \pm$ SD) |
| --- | --- | --- | --- |
| C1 | 4.75E-05 ± 7.54E-05 | 2.50E-05 ± 3.18E-05 | 8.35E-05 ± 5.14E-05 |
| C2 | 5.67E-05 ± 5.21E-05 | 4.34E-05 ± 6.29E-05 | 1.09E-04 ± 1.60E-04 |
| C3 | 7.64E-05 ± 8.39E-05 | 5.38E-05 ± 4.81E-05 | 8.85E-05 ± 9.70E-05 |
| C4 | 1.74E-04 ± 3.05E-04 | 6.80E-05 ± 4.40E-05 | 1.44E-04 ± 7.96E-05 |
| C5 | 2.74E-04 ± 1.74E-04 | 1.34E-04 ± 6.55E-05 | 2.03E-04 ± 1.15E-04 |
| C6 | 8.62E-04 ± 3.45E-04 | 1.53E-03 ± 3.05E-03 | 8.41E-04 ± 1.97E-04 |
| C7 | 1.60E-03 ± 2.81E-04 | 1.12E-03 ± 1.87E-04 | 1.44E-03 ± 5.29E-04 |
| C9 | 1.49E-02 ± 2.48E-03 | 1.09E-02 ± 1.73E-03 | 1.56E-02 ± 2.36E-03 |
| C10 | 9.83E-02 ± 3.93E-02 | 5.41E-02 ± 1.78E-02 | 7.68E-02 ± 1.16E-02 |
| C11 | 1.58E-01 ± 2.85E-02 | 1.17E-01 ± 8.84E-03 | 1.72E-01 ± 1.82E-02 |
| C12 | 3.82E-01 ± 1.32E-01 | 2.26E-01 ± 2.53E-02 | 3.28E-01 ± 3.77E-02 |
| C13 | 6.61E-01 ± 5.38E-02 | 4.47E-01 ± 7.78E-02 | 6.62E-01 ± 8.42E-02 |
| C14 | 1.30E+00 ± 1.19E-01 | 9.24E-01 ± 1.10E-01 | 1.38E+00 ± 1.38E-01 |

BPV-2
| Dilution | Naica ($\lambda \pm SD$) | QIAcuity ($\lambda \pm SD$) | LOAA ($\lambda \pm SD$) |
| --- | --- | --- | --- |
| C1 | 0.00 ± 0.00 | 2.88E-05 ± 3.12E-05 | 1.50E-04 ± 1.95E-04 |
| C2 | 3.74E-05 ± 4.18E-05 | 2.15E-05 ± 3.08E-05 | 3.63E-05 ± 3.65E-05 |
| C3 | 3.40E-05 ± 4.56E-05 | 3.94E-05 ± 4.43E-05 | 6.74E-05 ± 7.50E-05 |
| C4 | 8.55E-05 ± 6.51E-05 | 9.30E-05 ± 6.04E-05 | 1.56E-04 ± 1.15E-04 |
| C5 | 2.19E-04 ± 1.26E-04 | 1.60E-04 ± 7.79E-05 | 3.12E-04 ± 8.44E-05 |
| C6 | 9.15E-04 ± 2.50E-04 | 1.73E-03 ± 3.54E-03 | 1.02E-03 ± 3.25E-04 |
| C7 | 1.64E-03 ± 3.11E-04 | 1.57E-03 ± 2.45E-04 | 1.91E-03 ± 5.45E-04 |
| C9 | 1.72E-02 ± 2.31E-03 | 1.41E-02 ± 1.96E-03 | 1.87E-02 ± 3.52E-03 |
| C10 | 1.09E-01 ± 3.92E-02 | 6.90E-02 ± 2.23E-02 | 8.90E-02 ± 1.43E-02 |
| C11 | 1.73E-01 ± 3.25E-02 | 1.50E-01 ± 1.06E-02 | 2.05E-01 ± 2.03E-02 |
| C12 | 4.29E-01 ± 1.55E-01 | 2.86E-01 ± 2.70E-02 | 3.81E-01 ± 4.75E-02 |
| C13 | 7.52E-01 ± 5.11E-02 | 6.04E-01 ± 5.51E-02 | 7.72E-01 ± 9.77E-02 |
| C14 | 1.54E+00 ± 1.19E-01 | 1.20E+00 ± 1.10E-01 | 1.55E+00 ± 1.39E-01 |

IFN
| Dilution | Naica ($\lambda \pm \text{SD}$) | QIAcuity ($\lambda \pm \text{SD}$) | LOAA ($\lambda \pm \text{SD}$) |
| --- | --- | --- | --- |
| C1 | 1.25E-05 ± 2.49E-05 | 3.58E-05 ± 2.12E-05 | 3.02E-05 ± 4.50E-05 |
| C2 | 3.09E-05 ± 5.35E-05 | 5.37E-05 ± 4.20E-05 | 1.79E-05 ± 2.53E-05 |
| C3 | 5.29E-05 ± 8.04E-05 | 6.45E-05 ± 4.81E-05 | 8.01E-05 ± 7.05E-05 |
| C4 | 6.91E-05 ± 9.13E-05 | 1.16E-04 ± 4.32E-05 | 4.23E-05 ± 6.74E-05 |
| C5 | 1.31E-04 ± 8.26E-05 | 1.55E-04 ± 7.56E-05 | 1.21E-04 ± 5.77E-05 |
| C6 | 5.65E-04 ± 2.31E-04 | 1.50E-03 ± 2.66E-03 | 6.38E-04 ± 1.92E-04 |
| C7 | 1.31E-03 ± 2.83E-04 | 1.42E-03 ± 5.06E-04 | 9.65E-04 ± 4.91E-04 |
| C9 | 1.17E-02 ± 6.92E-04 | 1.13E-02 ± 1.49E-03 | 1.24E-02 ± 7.30E-04 |
| C10 | 6.92E-02 ± 2.56E-02 | 7.98E-02 ± 2.78E-02 | 6.17E-02 ± 6.56E-03 |
| C11 | 1.09E-01 ± 2.02E-02 | 1.87E-01 ± 1.93E-02 | 1.25E-01 ± 1.03E-02 |
| C12 | 2.73E-01 ± 1.03E-01 | 3.54E-01 ± 4.19E-02 | 2.70E-01 ± 2.61E-02 |
| C13 | 4.78E-01 ± 3.42E-02 | 7.57E-01 ± 6.52E-02 | 5.18E-01 ± 5.12E-02 |
| C14 | 9.88E-01 ± 6.47E-02 | 1.49E+00 ± 1.07E-01 | 1.06E+00 ± 7.70E-02 |

**Table S.5.** Average accepted partitions per reaction (mean ± SD) across platforms.

| Platform | Average partitions | SD |
| --- | --- | --- |
| LOAA | 18370.25 | 1230.10 |
| Naica | 15584.25 | 1349.49 |
| QIAcuity | 25169.71 | 1298.46 |

