## Supplementary Information for "Cross-platform digital PCR evaluation of bovine papilloma virus quantification: introducing PCR-ValiPal for Standardized guided assay validation"

Complete replicate datasets for all assays (BPV-1, BPV-2, IFN) across all platforms (Naica, QIAcuity, LOAA, and qPCR) are available in Supplementary Tables S6–S17 (Excel file).

Table S6. Complete replicate data for BPV-1 on the Naica platform.

Table S7. Complete replicate data for BPV-2 on the Naica platform.

Table S8. Complete replicate data for IFN on the Naica platform.

Table S9. Complete replicate data for BPV-1 on the QIAcuity platform.

Table S10. Complete replicate data for BPV-2 on the QIAcuity platform.

Table S11. Complete replicate data for IFN on the QIAcuity platform.

Table S12. Complete replicate data for BPV-1 on the LOAA platform.

Table S13. Complete replicate data for BPV-2 on the LOAA platform.

Table S14. Complete replicate data for IFN on the LOAA platform.

Table S15. Complete replicate data for BPV-1 from the CFX96 instrument.

Table S16. Complete replicate data for BPV-2 from the CFX96 instrument.

Table S17. Complete replicate data for IFN from the CFX96 instrument.
